# *De novo* macrocyclic peptide discovery produces potent class A GPCR ligands with diverse pharmacology

**DOI:** 10.64898/2026.10.21.760615

**Authors:** Fernando Salgado-Polo, Marin Matic, Philip Ben Rainsford, Felix Faas, Max Meyrath, Rajesh Regmi, Mette M Rosenkilde, Martyna Szpakowska, Andy Chevigné, Alexander S Hauser, Joseph M Rogers, Martin Gustavsson

**Affiliations:** Department of Biomedical Sciences, University of Copenhagen, Copenhagen, Denmark; Department of Drug Design and Pharmacology, Center for Pharmaceutical Data Science Education, University of Copenhagen, Copenhagen, Denmark; Immuno-Pharmacology and Interactomics, Department of Infection and Immunity, Luxembourg Institute of Health, Esch-sur-Alzette 4354, Luxembourg

**Keywords:** G protein-coupled receptor, Chemokine receptor, Antagonist, Drug discovery, mRNA display, Macrocyclic peptides, Structure prediction

## Abstract

G protein-coupled receptors (GPCRs) represent one of the largest protein families of therapeutic importance, as over one third of all currently approved drugs target GPCRs. A growing proportion of GPCR drugs are peptides. However, class A GPCRs (over 80% of clinically studied targets) are underexploited as peptide targets, presumably since their only structured domain is the core, seven-helix, integral membrane fold presenting a limited solvent-exposed surface area. Screening libraries of *de novo* macrocyclic peptides have allowed for the targeting of otherwise intractable proteins, but their potential to generate selective ligands for class A GPCRs has not been demonstrated. Here, we used random nonstandard peptides integrated discovery (RaPID) to identify *de novo* macrocyclic peptide ligands targeting the chemokine receptor CXCR4. CXCR4 regulates cell migration, immune surveillance, and inflammation, making it an important therapeutic target for multiple diseases. We discovered novel, potent CXCR4-selective antagonists binding within the orthosteric pocket with nanomolar affinity. The most active peptides effectively inhibited receptor signaling, suppressed chemokine-driven cell migration, and blocked CXCR4-driven HIV entry. Our results show that even the structurally minimal class A GPCRs are viable targets for *de novo* cyclic peptide discovery, highlighting the potential for uncovering ligands towards other members of the GPCR superfamily.

## Main

The superfamily of G protein-coupled receptors (GPCRs) contains over 800 membrane proteins divided into six classes (A-F) based on sequence and functional similarity. They all recognize extracellular stimuli, triggering coupling of intracellular transducers, such as G proteins and arrestins, in a myriad of pathophysiological processes^1^. Because of this, GPCRs form one of the largest families of therapeutic targets, with ∼36% of all approved drugs targeting them^2^. Most drugs targeting GPCRs are small molecules but, peptides have emerged as a promising drug modality for GPCR targeting, with several recent successful therapies targeting class B GPCRs. With high target selectivity and favorable safety profiles, peptides offer several potential benefits over small-molecule drugs^3^.

The large extracellular domains present in class B receptors, such as glucagon-like peptide-1 receptor (GLP1R), present a large interaction surface with their ligands and make them beneficial targets for peptide therapies. However, most clinically studied GPCR targets (∼86%) are class A GPCRs^4,5^. Relative to class B, class A receptors have a limited solvent-exposed area for ligand recognition, restricted to a short N-terminal region, three extracellular loops (ECLs), an orthosteric pocket, and allosteric binding sites. Thus, class A GPCRs could pose significant challenges for peptide-based drug discovery, and these receptors remain an underexploited source of peptide-based therapeutic opportunities.

Macrocyclic peptides often show higher selectivity and affinity, due to their restricted peptide conformations^6^. Random nonstandard peptides integrated discovery (RaPID) has emerged as a leading technique in the discovery of macrocyclic peptides^7^. RaPID combines mRNA-display technology with ribosomally synthesized thioether-cyclized peptides generated by genetic code reprogramming ^8,9^. This allows for the *in vitro* screening of libraries containing over ten trillion unique macrocyclic peptides. In the past decade, the RaPID system has generated high-affinity, selective ligands for numerous previously intractable targets, including several enzymes and soluble proteins^10–13^, as well as membrane transporters solubilized in detergent micelles, such as the MATE family, P-glycoprotein, and NTCP to block hepatitis B viral entry^14–16^. Recently, RaPID has been used to target the bacterial receptor histidine kinase AgrC that was reconstituted in nanodiscs^17^. Nanodiscs are soluble lipid bilayers mimicking biological membranes which are preferred to identify receptor ligands in defined native-like lipid environments^18,19^ Overall, suggesting that RaPID may be a powerful and unexploited drug-discovery tool applicable to many class A GPCRs.

Among class A receptors, the group of over twenty chemokine receptors have emerged as promising therapeutic targets, as they regulate cell migration, immune surveillance, and inflammation in response to over 40 small (8-12 kDa) proteins known as chemokines ^20^. Dysregulation in this system has been implicated in a wide range of diseases, including cancer, autoimmune disorders, and viral infections^21^. Five drugs targeting chemokine receptors have been approved by the FDA^2^. First, the small-molecule CCR5 antagonist maraviroc (selzentry) was approved for the treatment of HIV-1 infections^22^, followed by the small-molecule CXCR4 antagonist plerixafor (mozobil, AMD3100) for hematopoietic stem cell mobilization^23,24^, and the anti-CCR4 monoclonal antibody mogamulizumab-kpkc (poteligeo) against mycosis fungoides and Sézary syndrome^25^. More recently, two more drugs targeting CXCR4 have been approved: the small-molecule antagonists mavorixafor (xolremdi) for the treatment of WHIM syndrome^26^, and the poly-cationic, natural product derivative^27^, disulfide-bridged peptide motixafortide (aphexda) for the same indication as plerixafor^28^. As such, CXCR4 is the archetypical chemokine receptor, and the ideal target to evaluate *de novo* cyclic peptide screening.

Here, we established a high-throughput screening pipeline where RaPID was adapted for the first time to target a class A GPCR, to discover *de novo* macrocyclic peptides against the chemokine receptor CXCR4. Targeting nanodisc-reconstituted CXCR4, we identified novel, potent, and functionally diverse *de novo* macrocyclic peptide binders, which modulate the receptor through distinct mechanisms of action. Our findings demonstrate the potential of the RaPID system to uncover pharmacologically diverse ligands for challenging GPCR targets and provide a foundation for precise modulation of chemokine signaling.

## Results

### RaPID identifies novel CXCR4-binding peptides

RaPID is a highly customizable system that allows for the introduction of unnatural amino acids in peptide screening libraries. Using genetic code reprogramming of the ATG codon, the N-terminal amino acid did not encode a methionine but a chloroacetyl-D-phenylalanine. Upon translation, this residue reacted with the thiol of an encoded cysteine, cyclizing the peptide by a non-reducible thioether bond.

For the RaPID selections (**Fig.1a**), we purified wildtype human CXCR4, which was reconstituted in MSP1D1 nanodiscs containing 1-palmitoyl-2-oleoyl-*sn*-glycero-3-phosphocholine (POPC) and cholesterol (9:1, w/w)^29^. To exclude non-specific lipid binders, the library underwent “negative” selections in each round (apart from the first), by depleting the library by pull-down with immobilized empty biotinylated MSP1D1 nanodiscs (**Fig.1a**). The “positive” selections were then performed in one of two modes: immobilizing the nanodiscs using biotinylated MSP1D1 (Screen 1, MSP-immobilized) or directly capturing C-terminally biotinylated CXCR4 (Screen 2, receptor-immobilized). At the end of each selection round, a qPCR was performed to determine total library recovery in the negative and positive selections, as well as the recovery ratio between them (**Fig.S1A**). The results of Screen 1 indicated that positive peptide recovery never surpassed that of the negative selection, which could be due to a presence of both empty and receptor nanodiscs in the positive selection. In contrast, direct receptor immobilization in Screen 2 achieved a 100-fold larger positive recovery.

Next-generation sequencing (NGS) revealed tens of peptides enriched for CXCR4 over empty nanodiscs (**Fig.1b**). In the cases where several CXCR4-targeting peptides showed near-identical sequences, only the most enriched candidate was kept for further analysis. Thus, 11 peptides with unique sequences were selected, named alphabetically **A**-**K** (**Fig.1c**, **Table S1**). The peptide candidates showed a high arginine content, but they did not present strong sequence identity (**Fig.S1b, Table S1**). They also differed in cycle size and net charge, where **A** and **E** were the only linear and zwitterionic peptides, respectively (**Fig.S1c**, **Table S1**). The identified peptides did not show sequence similarity to the FDA-approved cyclic peptide motixafortide and were notably less charged (**Fig.S1b-c**, **Table S1**). All peptides (except for **J**) could be synthesized at mg scale.

The peptide candidates were selected to be CXCR4 binders with no bias to interact with a specific region of the receptor. Because of this, they could bind to the orthosteric pocket, intracellularly or allosterically. Therefore, we next evaluated the ability of the peptides to displace fluorescently labeled CXCL12 from the extracellular side of CXCR4 using fluorescence polarization (**Fig.1d**, **S1d**). The results showed that peptides **F**, **G**, **H** and **I** significantly reduced CXCL12 binding to CXCR4 (**Fig.1e**, **S1e**). Importantly, even though the small-molecule inverse agonist IT1t^30^ binds in the orthosteric pocket^31^, it did not show a decrease CXCL12 binding. This indicates that macrocyclic peptides are superior to IT1t in causing complete displacement of CXCL12 from high concentrations of CXCR4 (**Fig.1e**, **S1e**).

To gain insight into the ability of the identified peptides to prevent ligand-induced receptor activation, we then used a fluorescence saturation binding assay (FSBA) to detect their effect on direct G protein binding (**Fig.1f**). For this, we employed mini-Gα_o_, which is efficiently recruited to CXCR4 in the presence of CXCL12 compared to the apo receptor (**Fig.1g**). Addition of the identified peptides to ligand-bound CXCR4 showed different effects, where peptides **G** and **I** significantly decreased *in vitro* mini-Gα_o_ recruitment, and peptides **F** and **K** showed partial reduction (**Fig.1g**). The strong binder IT1t was able to outcompete CXCL12-driven receptor activation, similar to the reduction achieved with peptide **I**. Thus, our biophysical experiments support that the identified peptides compete with CXCL12 to bind at the extracellular side of the receptor and/or antagonize intracellular G protein binding.

**Fig. 1:**
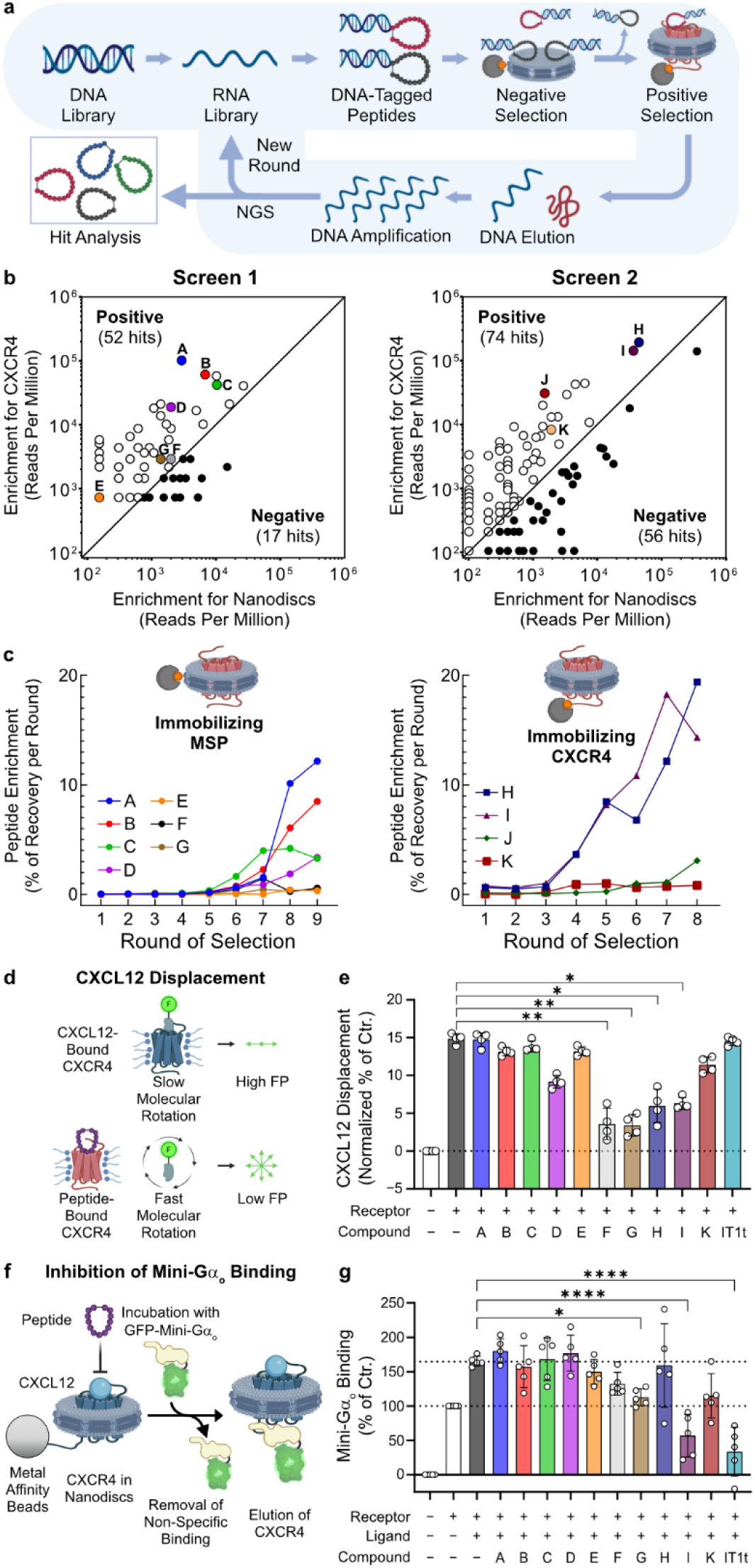
Discovery of novel peptides targeting CXCR4 using RaPID. **a**, RaPID workflow for the identification of CXCR4-specific peptides. **b**, Enrichment plots showing all peptides detected in the last round from Screen 1 (left) and Screen 2 (right). Peptides that were more abundant in the positive (binding CXCR4) or negative (binding empty nanodiscs) selections are plotted above or below the diagonal, respectively. **c**, Peptide enrichment over rounds of RaPID selection for the two screens. The enrichment of each peptide refers to its abundance with respect to total peptide recovery per round. **d-e**, Fluorescence polarization assay detecting the displacement of 12.5 nM CXCL12-AF488 from 500 nM CXCR4 by the identified peptides (5 µM). (**d**) Schematic representation of the assay; (**e**) measured changes in fluorescence polarization in the presence of 5 µM peptides or IT1t. **f-g**, *in vitro* fluorescent saturation binding assay (FSBA) using C-terminally His10-tagged CXCR4 in MSP1D1 nanodiscs and N-terminally GFP-tagged mini-Gαo. (**f**) Schematic depiction of the assay; (**g**) effect of CXCL12 (0.5 µM), the novel peptides (5 µM), or IT1t (5 µM) on receptor interaction with mini-Gαo. (**a**,**d**,**f**) Created with BioRender.com. (**e**,**g**) The asterisk symbols indicate statistically significant differences (*p < 0.05, **p < 0.01, ****p < 0.0001) between the indicated groups, determined by one-way ANOVA. Only significant differences to the control group are shown. N = 3 independent experiments, performed in technical triplicates. All data are shown as mean ± SD.

### Novel peptides are strong CXCR4 inverse agonists

We next examined the role of the identified peptides in cell signaling. After observing that the viability of HEK293A cells was not affected upon incubation with 10 µM of the identified peptides (**Fig.S1e**), we determined the peptides’ ability to modulate mini-Gα_s/i_ coupling by bioluminescence resonance energy transfer (BRET)-based assays (**Fig.2a**). While addition of CXCL12 induced CXCR4 activation and subsequent mini-Gα_s/i_ coupling, none of the identified peptides displayed such agonistic functions (**Fig.S2a**).

Performing the assay in the presence of 10 nM CXCL12 indicated that five peptides acted as antagonists with differing effects on receptor efficacy (E_max_) and peptide antagonistic potency (IC_50_) (**Fig.2b-c**, **S2b**). While peptides **A**-**E** did not affect mini-Gα_s/i_ coupling, peptides **F** (IC_50_ = 604 nM) and **G** (IC_50_ = 50 nM) showed complete and partial antagonism, respectively. In contrast, peptides **H** (IC_50_ = 150 nM), **I** (IC_50_ = 23 nM), and **K** (IC_50_ = 29 nM) were high-potency full antagonists, whose potency closely resembled that of IT1t (IC_50_ = 46 nM) (**Fig.2b-c**, **S2b**).

Since downstream responses are triggered upon G protein heterotrimer dissociation, we further determined the peptides’ function measuring BRET changes upon recruitment of fluorescently labeled Gβγ to nanoluciferase (Nluc) fused to the C-terminal tail of GRK3^32^ (**Fig.2d**). Although the results largely agreed with those seen with mini-Gα_s/i_, peptide **F**, but not **G**, acted as a receptor antagonist (IC_50_ = 662 nM) (**Fig.2e-f**, **S2c**). The heterotrimer dissociation assay allows for the detection of changes below constitutive activity (E_min_) levels, which were observed for the inverse agonist IT1t (IC_50_ = 522 nM). Similarly, peptides **H** (IC_50_ = 2.6 µM), **I** (IC_50_ = 200 nM), and **K** (IC_50_ = 1.2 µM) behaved as inverse agonists, thus outcompeting CXCL12 and establishing interactions that reduce CXCR4 basal activity^33^ .

Receptor activation triggers phosphorylation by G protein-coupled receptor kinases (GRKs) and subsequent β-arrestin recruitment, which we followed using a BRET-based setup (**Fig.2g**). CXCL12-induced β-arrestin2 recruitment was blocked with micromolar potencies by peptides **D**, **E**, and **F**, while peptide **G** showed a potency of 119 nM (**Fig.2h-i**, **S2d**). In contrast, peptides **H** (IC_50_ = 133 nM), **I** (IC_50_ = 54 nM), and **K** (IC_50_ = 120 nM) showed nanomolar potencies as β-arrestin2 antagonists (**Fig.2h-i**, **S2d**). Interestingly, this is the only assay where our peptides did not outperform IT1t (IC_50_ = 8 nM) as β-arrestin2 antagonists.

GRK phosphorylation also leads to receptor internalization, which can be detected using a diffusion-enhanced resonance energy transfer (DERET) assay (**Fig.2j**). Only peptides H, **I**, and, **K** significantly reduced CXCL12-induced CXCR4 internalization down to the basal levels observed in unstimulated cells, and similar to the antagonistic effect of IT1t (**Fig.2k-l**, **S2e**).

Overall, seven of the *de novo* peptides identified by RaPID behaved as CXCR4 antagonists, showing similar pharmacological properties in the direct recruitment of mini-G and β-arrestin2, as well as G protein activation (**Fig.2m**). While **I** and **K** were the most potent peptides in both readouts, peptide **G** was able to fully antagonize β-arrestin2 recruitment but only partially reduced mini-Gα_s/i_ coupling and had no effect on G protein activation even at the highest concentrations tested, showing selectivity to inhibit one of the two pathways (**Fig.2m**).

### Novel peptides suppress CXCR4-dependent migration and viral fusion

In a physiological context, CXCR4 downstream signaling leads to cell migration towards CXCL12 gradients. We therefore evaluated the ability of peptides **H, I** and **K** to block motility using the immortalized T Jurkat cell line in trans-well migration chambers. Our results showed that all three peptides were able to suppress migration toward CXCL12 as efficiently as the small-molecule inverse agonist IT1t **Fig.2n**).

CXCR4 has also been implicated in HIV entry, where it acts as a co-receptor to CD4 in the binding and fusion of viral particles expressing gp120 (**Fig.2o**). Using a luminescence-based approach in HEK293 cells, we allowed effector cells to fuse with CXCR4-expressing target cells (**Fig.2p**). Addition of 10 µM of peptide **I** reduced viral fusion events with similar efficiency to IT1t. Overall, our results provide translational evidence for peptide **I** to block CXCR4-mediated functional cellular responses.

**Fig. 2:**
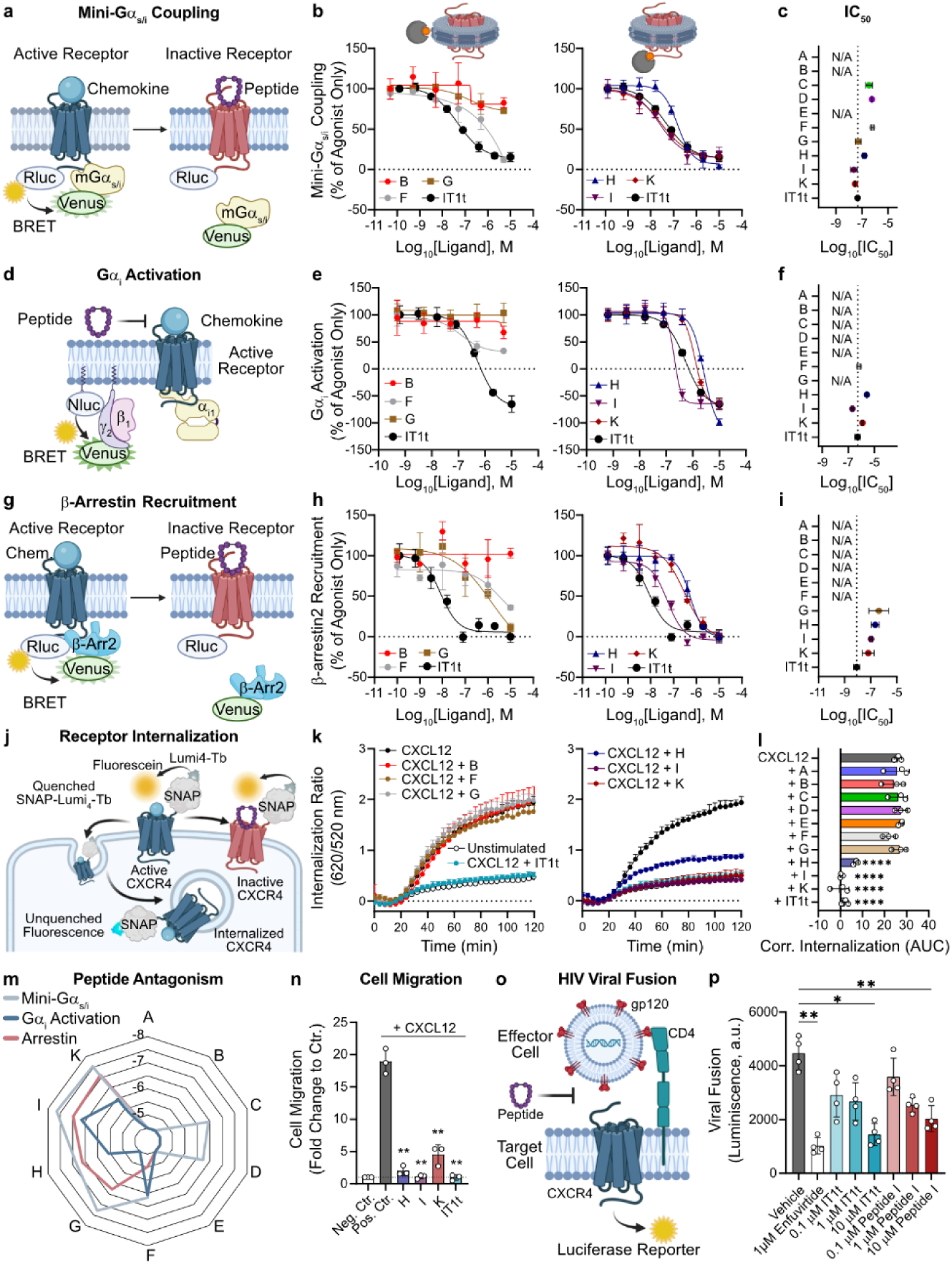
Peptides identified by RaPID antagonize CXCR4-mediated cellular functions. **a**,**b**,**c**, Antagonistic effects of the identified peptides on direct mini-Gαs/i coupling to CXCR4. (**a**) Assay scheme for the bioluminescence resonance energy transfer (BRET)-based detection of mini-Gαs/i coupling to CXCR4 in response to CXCL12 and antagonized by the identified peptides. Cells were stimulated with CXCL12 (10 nM) after incubation with each peptide at different concentrations (**b**), allowing for the quantification of their antagonistic potencies (**c**). **d**,**e**,**f**, Function of the novel peptides on CXCR4-driven Gαi activation. (**d**) Schematic depiction of the Gαi activation assay employed to determine the antagonistic function of the identified peptides. BRET signal was detected upon the translocation of the β1 and venus-tagged γ2 subunits G protein subunits to the plasma membrane, where can come into proximity to the NanoLuc® luciferase (Nluc)-tagged C-terminal tail of GRK3. Peptide activity is shown as dose-response curves upon addition of 10 nM CXCL12 (**e**), and the potency of each of them (**f**). **g**,**h**,**i**, Changes in direct β–arrestin2 recruitment by novel CXCR4-targeting peptides. (**g**) Scheme of the direct recruitment of N-terminally venus–tagged β–arrestin2 to C–terminally Rluc3–tagged CXCR4 detected by BRET. Peptides were initially incubated with CXCR4 and allowed to be displaced by CXCL12. Dose-response curves for each peptide (**h**), and their antagonistic potency (**i**). **j**,**k**,**l**, Diffusion-enhanced resonance energy transfer (DERET)-based CXCR4 internalization assay. (**j**) Schematic representation of the internalization assay, where surface N-terminally SNAP-tagged CXCR4 was labeled with Lumi4-Tb. While on the cell surface, light emitted by the fluorophore is absorbed by cell-impermeable fluorescein (quencher); receptor internalization leads to increased signal due to lower intracellular fluorescein concentration. The antagonistic effect of the identified peptides (5 µM) on CXCL12-induced internalization was compared to that of IT1t (5 µM) and CXCL12 alone (100 nM). Data is shown as time-course changes in the internalization ratio (**k**), and the area under the curve for each ligand (**l**). **m**, Radar plot showing the antagonistic potency of each peptide toward the direct recruitment of mini-Gαs/I or β–arrestin2 to CXCR4, and Gαi protein activation. **n**, Effect of the identified peptides (1 µM) or IT1t (1 µM) blocking trans-well migration of Jurkat cells in response to 10 nM CXCL12. **o**,**p**, Effect of peptide **I** and IT1t on CXCR4-mediated HIV viral fusion. (**o**) Scheme of the cell fusion assay driven by the interaction between gp120 and CD4, (**p**) where the influence of CXCR4 antagonists was measured and compared to vehicle; enfuvirtide was used as a fusion inhibitor. (**a**,**d**,**g**,**j**,**o**) Created with BioRender.com. (**l**,**n**,**p**) The asterisk symbols indicate statistically significant differences (*p < 0.05, **p < 0.01, ****p < 0.0001) between the indicated groups, determined by one-way ANOVA. Only significant differences to the control group are shown. (**b-c**,**e-f**,**h-I**,**k-l,n**,**p**) N = 3 independent experiments, performed in technical triplicates. All data are shown as mean ± SD.

### Peptide I shows high selectivity toward CXCR4

We next determined whether the identified peptides are selective for CXCR4. Due to the structural similarity between CXCR4 and ACKR3 and the activation-prone nature of ACKR3^34^, CXCR4 antagonists can frequently function as ACKR3 agonists, such as the FDA-approved drug Plerixafor (AMD3100)^24^. Thus, we determined the extent to which the peptides activate ACKR3. ACKR3 cannot couple to G proteins, but it can recruit β-arrestin2 in response to full agonists like VUF11207^35^, which was validated to have an agonistic potency (EC_50_) of 8 nM in a BRET-based assay (**Fig.3a-b**, **S3a**). Stimulation with our novel peptides showed that seven of them were full ACKR3 agonists. Even though **B** and **C** did not show any activity toward CXCR4, they were selective ACKR3 agonists, with EC_50_ values of 30 nM and 824 nM, respectively (**Fig.3a-b**, **S3a**). While peptide **H** acted as the most active identified peptide toward ACKR3 (EC_50_ = 22 nM), the potency of **I** for ACKR3 (EC_50_ = 890 nM) was 15-fold lower compared to its potency for CXCR4 (**Fig.3a-b**, **S3a**). Lastly, peptide **K** was the most selective peptide for CXCR4, as it only triggered 20% β-arrestin2 recruitment to ACKR3 at a concentration of 10 µM.

In contrast to CXCR4, ACKR3 undergoes high constitutive internalization and recycling, resulting in only a ≈ 50% increase in the DERET assay upon stimulation by VUF11207 or CXCL12, one of the endogenous ligands of ACKR3 (**Fig.3c-d**). The results showed that peptides **B**, **F**, **G**, and **H** promoted ACKR3 internalization (**Fig.3c-d**, **S3b**). This assay also confirmed that **I** and **K**, which were both strong antagonists of CXCR4 (**Fig.2k-l**), had a weaker effect on ACKR3 activity (**Fig.3c-d**).

To further examine the selectivity of our peptides, we next compared their ability to recruit β-arrestin across different receptors. Our prior results showed that most peptides had similar potencies for CXCR4 and ACKR3, except for peptides **B**, selectively targeting ACKR3, and peptides **I** and **K**, preferring CXCR4 (**Fig.3e**). As peptide **I** displayed higher potency, we employed it at a fixed concentration of 5 µM to stimulate all chemokine receptors, each of them overexpressed separately in HEK293 cells (**Fig.3f**). Analysis of potential agonistic roles of peptide **I** confirmed that it promotes ≈ 60% increase in arrestin recruitment to ACKR3 at 5 µM. As an antagonist, peptide **I** selectively suppressed over 85% of CXCR4 activation, with no off-target activity larger than 5% for any other chemokine receptor (**Fig.3f**, **Table S3**).

We then investigated the ability of peptide **I** to displace fluorescently labeled CXCL12 in a cellular environment (**Fig.3g**). Ligand displacement in cells overexpressing CXCR4 showed that binding of peptide **I** (IC_50_ = 26 nM) was significantly stronger compared to unlabeled CXCL12 (IC_50_ = 46 nM), whose apparent affinity resembled that of the FDA-approved drug AMD3100 (IC_50_ = 58 nM) (**Fig.3h**). CXCL12 can also be recognized by the atypical receptors ACKR3 and ACKR5, which is a constitutively internalizing receptor with high, agonist-independent association with β-arrestin^36,37^. Therefore, we tested how strongly peptide **I** would bind to the two other CXCL12 receptors. While peptide **I** (IC_50_ = 4 µM) showed a 2,000-fold lower apparent affinity than CXCL12 (IC_50_ = 2 nM) for ACKR3 (**Fig.3i**), the binding of peptide **I** to ACKR5 (IC_50_ = 325 nM) was closer to that of CXCL12 (IC_50_ = 131 nM) (**Fig.3j**, **S3c**).

Overall, our results indicate that peptide **I** selectively antagonizes CXCR4 at its IC_90_ (58 nM), where minimal off-target activity would be anticipated toward closely related receptors (**Fig.2h**, **3a**,**f**,**h-j**). Moreover, this shows that our RaPID screens yielded peptides with multi-receptor selectivity.

**Fig. 3:**
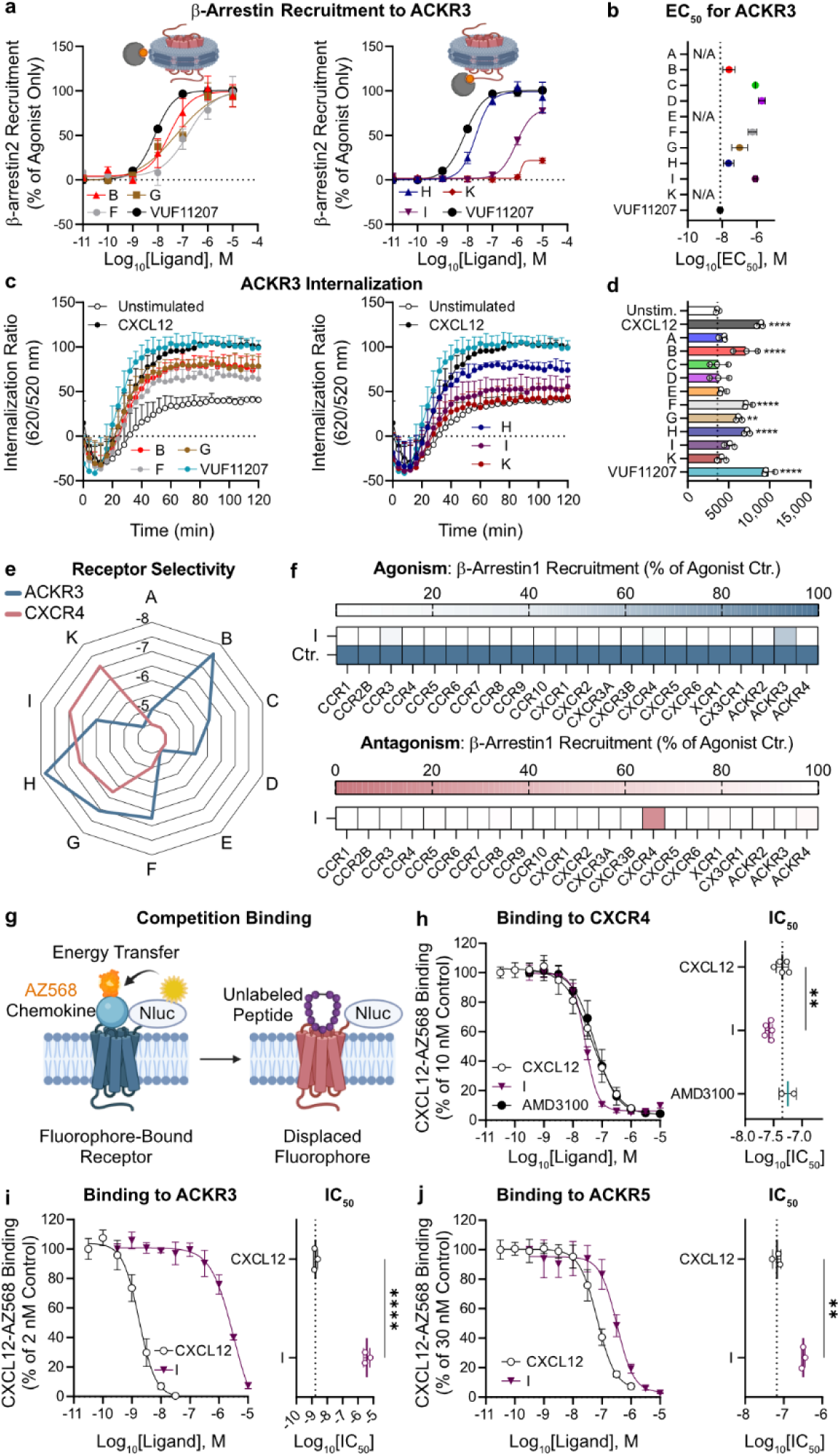
Novel antagonists show differential selectivity for CXCR4 over other chemokine receptors. **a**,**b**, Agonistic effect of the peptides identified by RaPID on β-arrestin2 recruitment to ACKR3 using a direct BRET-based assay. The results are shown as dose-response curves for each peptide (**a**), and the quantification of their potency (**b**). The full agonist VUF11207 was used as positive control for ACKR3 activation. **c**,**d**, Induction of ACKR3 internalization by the identified peptides (1 µM), compared to CXCL12 (100 nM) and VUF11207 (1 µM). Data is shown as time-course changes in the internalization ratio (**c**), and the area under the curve for each ligand (**d**). **e**, Summarizing radar plot displaying the differential selectivity of each peptide for the structurally related receptors ACKR3 and CXCR4. **f**, Selectivity screen for the effects of peptide **I** (5 µM) across all chemokine receptors. NanoLuciferase complementation-based assays with HEK293 cells overexpressing each receptor were performed to detect the agonism (top panel) or antagonism (bottom panel) of peptide **I** on the direct recruitment of β-arrestin1. Since ACKR1 does not recruit arrestins, and ACKR5 does not present ligand-induced arrestin recruitment, antagonism of peptide **I** could not be evaluated for them. **g**,**h**,**i**,**j**, BRET-based binding assay in HEK293 cells based on the competition of fluorescent CXCL12-AZ568 by other ligands. (**g**) Assay scheme (Created with BioRender.com). Displacement of labeled CXCL12 by unlabeled CXCL12 or peptide **I** from N-terminally Nluc-tagged CXCR4 (**h**), ACKR3 (**i**), or ACKR5 (**j**). The results are shown as dose-response curves (left) and the potencies for each ligand to bind each receptor. (**d**,**h-j**) The asterisk symbols indicate statistically significant differences (**p < 0.01, ****p < 0.0001) between the indicated groups, determined by one-way ANOVA. Only significant differences to the control group are shown. (**a-d**,**h-j**) N = 3 independent experiments, performed in technical triplicates. All data are shown as mean ± SD.

### Novel peptides specifically interact in the orthosteric pocket of CXCR4

The pharmacological characterization of peptide **I** showed its function as a potent inverse agonist for CXCR4. Our results also showed that this peptide competes for binding with CXCL12 (**Fig.1e**,**3h**); however, understanding its mechanism requires structural insights. Thus, we first detected which residues in peptide **I** are involved in interactions with nanodisc-embedded CXCR4 using water-ligand observed via gradient spectroscopy (waterLOGSY) by nuclear magnetic resonance (NMR) (**Fig.4a-c**). Peptide **I** spectra showed weak positive waterLOGSY signals for non-exchangeable protons in buffer, and stronger positive waterLOGSY signals when incubated with both empty and CXCR4 nanodiscs. This indicated that peptide **I** can interact with both empty and CXCR4-containing nanodiscs. Specific protons of peptide **I** that could be unambiguously assigned to the Hζ^3^ & Hη^2^ positions of Trp9, and the CH_3_^δ^ of Leu6 and Leu10 (**Table S4**) allowed us to compare their change in solvent exposure upon CXCR4 binding (**Fig.4b-c**). In the presence of CXCR4, Hζ^3^ & Hη^2^ in Trp9 were more solvent-exposed than in empty nanodiscs, *i.e.* lower waterLOGSY signal. In addition, when bound to CXCR4, one of the CH_3_^δ^ of Leu6 was more solvent-exposed than those of Leu10 (**Fig.4b-c**). These results confirm that peptide **I** binds to CXCR4 in a distinct pose, influencing the chemical environment of Leu6 and Trp9 in the peptide.

The interaction between CXCR4 and CXCL12 has been thoroughly studied, and key interacting receptor residues have been reported^38,39^. To better understand what residues may participate in binding peptide **I**, we generated alanine mutations of four CXCR4 amino acids interacting with CXCL12 where activation was not abrogated: W94^2.60^, D187^ECL2^, D262^6.58^, and H281^7.32^ (residue numbering follows the Ballesteros/Weinstein scheme) (**Fig.S5a**). CXCL12 showed a reduced potency to induce mini-Gα_s/i_ coupling to CXCR4 when W94^2.60^ or D187^ECL2^ were mutated to an alanine. We next investigated the effect of the same CXCR4 mutants on the potency of peptide **I** (**Fig.4d**). From the four residues, only mutation of D187^ECL2^, located in ECL2 with its side chain facing the orthosteric pocket, weakened the potency of peptide **I** for CXCR4. This indicates that peptide **I** interacts with ECL2 but binds the receptor in a distinct manner to that of CXCL12.

To complement our mutagenesis analysis, we used AlphaFold 3 co-folding to jointly predict structures of CXCR4 in complex with peptide **I**, peptide **H**, or CXCL12^40,41^. The predicted complexes were refined using Rosetta FastRelax^42^, and receptor-ligand contacts were identified using PLIP^43^ to compare shared and ligand-specific receptor contact patterns (**Fig.4e-h**). The model showed that peptide **I** formed ten unique interactions with CXCR4 residues, including six hydrogen bonds and salt bridges, namely Y121^3.37^, Q200^5.39^, Q202^5.41^, Y255^6.51^, H281^7.32^, Y190^ECL2^ (**Fig.4e-f,h, S5b**). Moreover, peptide **I** and CXCL12 had nine interaction sites in common, containing six hydrogen bonds and salt bridges: K37^1.31^, D97^2.63^, D262^6.58^, E277^7.28^, E288^7.39^, D187^ECL2^ (**Fig.4e-f,h, S5b**). Importantly, while peptide **I** contacted W94^2.60^ by aromatic π stacking, only mutation of the salt bridge with D187^ECL2^ compromised inhibition of CXCR4 (**Fig.4d**). Prediction of the binding mode of peptide **H** to CXCR4 showed eight interactions that were not shared with CXCL12, including seven unique hydrogen bonds, and it interacted with nine residues also recognized by CXCL12 (**Fig.4e,g-h, S5b-c**). Overall, CXCR4 is predicted to involve ∼5 more residues in interactions with peptide **I** compared to **H** and CXCL12 (**Fig.4e**). In agreement with the NMR results, the solvent exposed Leu6 of peptide **I** points away from the binding pocket, while the more buried Leu10 makes a hydrophobic contact with CXCR4.

To gain more insight into receptor preference, we investigated the mode of binding of peptide **H** to ACKR3. We first produced alanine mutants of four previously reported ACKR3 residues interacting with CXCL12^44^: W100^2.60^, E213^5.39^, Y268^6.51^, and Q301^7.39^ (**Fig.S5d**). The results showed a reduction in CXCL12 potency for the activation β-arrestin2 recruitment to ACKR3 upon mutation of W100^2.60^, but not when E213^5.39^ was mutated to an alanine. As previously described, mutation of Y268^6.51^, or Q301^7.39^ abrogated CXCL12-driven ACKR3 activation (**Fig.S5d**). Using the same ACKR3 mutants, we analyzed their effect on the potency of peptide **H**. From these four residues, mutants of Y268^6.51^, or Q301^7.39^ both abrogated receptor activity, while mutation of E213^5.39^, but not W100^2.60^, reduced the potency of peptide **H** for ACKR3 (**Fig.S5e**). This suggests that **H** has a mode of binding that partially differs from that of CXCL12.

**Fig. 4:**
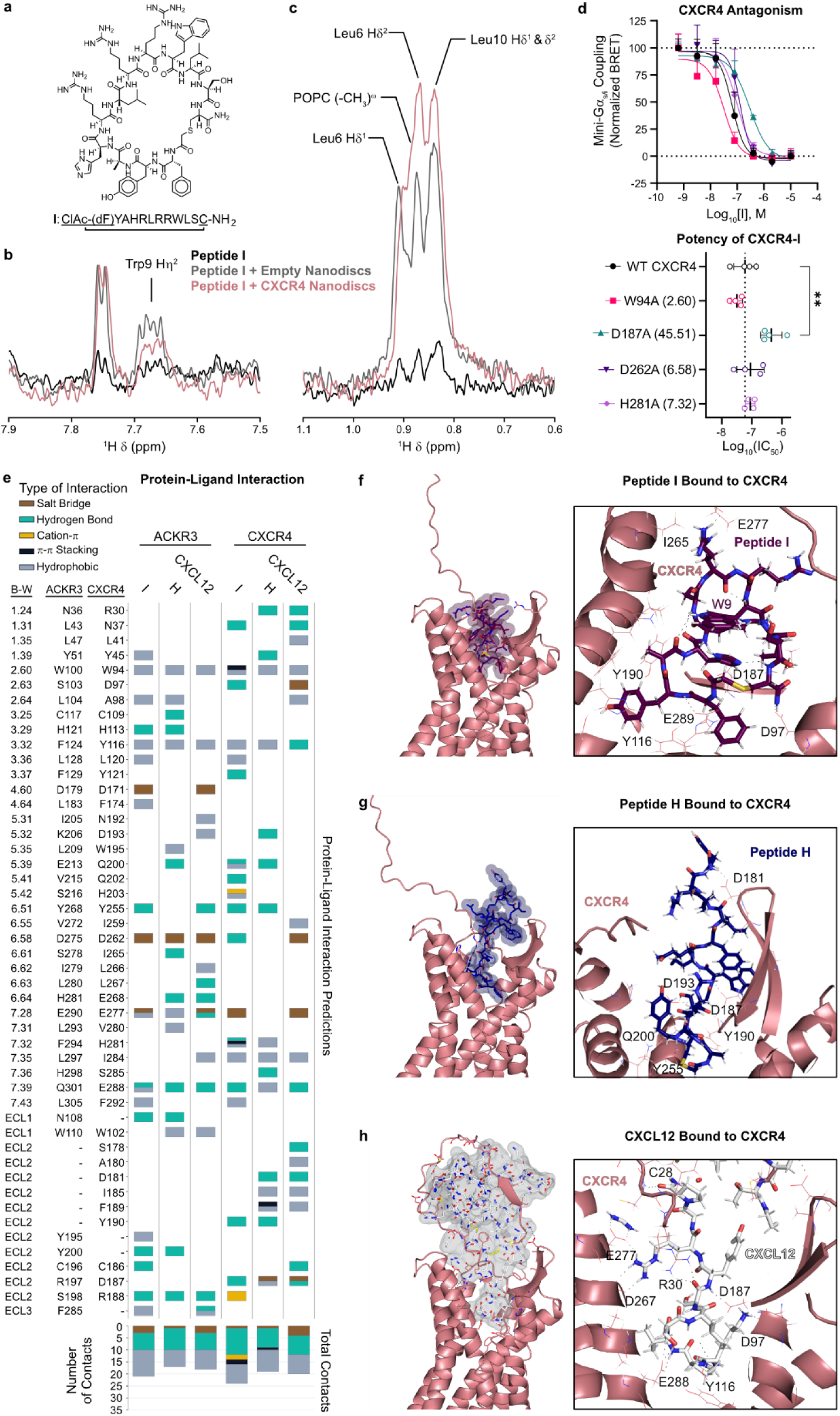
The most potent CXCR4 inverse agonists specifically interact in the orthosteric pocket. **a**, Structure and sequence of peptide **I**, showing the cyclization site between the N-terminal ClAc-(dF) and Cys12. **b,c**, Highlighed intensity changes from waterLOGSY spectra, specifically for (**b**) Trp^9^ and (**c**) Leu^6^ and Leu^10^. Spectra show peptide **I** in buffer (black), with empty POPC nanodiscs (grey), and CXCR4-containing POPC nanodiscs (red). The potential overlap between the signals from the CH3 groups in Leu^6^ and the lipid fatty acid tail was considered in the assignment. Full spectra are provided in Fig.S4. **d**, BRET-based assay detecting peptide **I**-mediated antagonism of mini-Gαs/i coupling to CXCR4 mutants. A fixed concentration of 10 nM CXCL12 was used to stimulate cells upon incubation with peptide **I**. The results are shown as dose-response curves (left), and the quantification of the antagonistic potency of peptide **I** for each mutant. **e**, Interaction matrix classifying the predicted contacts with ACKR3 and CXCR4 residues into five interaction categories. The Ballesteros/Weinstein nomenclature is followed; ECL amino acids without conservation are referred to using their residue number in each receptor. **f**,**g**,**h**, Predicted mode of binding between CXCR4 (in pale pink) and peptide **I** (**f**, in purple), peptide **H** (**g**, in dark blue), or CXCL12 (**h**, in gray). (Left) View of the extracellular site of CXCR4 (in ribbon representation), and the van der Waals radius of each ligand (in surface representation). (Right) Residues at a maximum distance of 2.5 Å from each ligand, including hydrogen bonds. Images created with PyMOL. (**d**) The asterisk symbols indicate statistically significant differences (**p < 0.01) between the indicated groups, determined by one-way ANOVA. Only significant differences to the control group are shown. N = 3 independent experiments, performed in technical triplicates. All data are shown as mean ± SD.

Next, we applied the same co-folding, relaxation, and contact-analysis workflow to ACKR3 complexes with peptides **I**, **H**, and CXCL12 (**Fig.S5f-g**). Our models for **H** showed nine unique interactions that were not present for CXCL12, containing six hydrogen bonds and salt bridges: C117^3.25^, H121^3.29^, E213^5.39^, S278^6.61^, N108^ECL1^, and Y200^ECL2^ (**Fig.4e, S5b,g-h**). This confirmed our results involving Glu^5.39^ as a probable interactor for peptide **H** (**Fig.S5e**). Importantly, peptide **H** was predicted to bind in a conformation that is vertically mirrored to its binding mode to CXCR4. Because of this, **H** had contacts with residues that were not involved in ligand binding in any of the other models: including the aforementioned hydrogen bonds with C117^3.25^, S278^6.61^, as well as the hydrophobic interaction with L293^7.31^. Moreover, we also identified eight shared ACKR3 residues interacting with both **H** and CXCL12, including the salt bridge with D275^6.58^, and the hydrogen bonds with H281^6.64^, Q301^7.39^, and S198^ECL2^ (**Fig.4e, S5b,g-h**). Lastly, we predicted the binding mode of peptide **I** to ACKR3, showing ten unique interactions that were not shared with CXCL12, and nine residues that were also recognized by CXCL12 (**Fig.4e, S5b,f,h**). In summary, although CXCL12 had a similar number of total interactions with CXCR4 or ACKR3, both peptides **H** and **I** showed a higher number of total and specific contacts with CXCR4 than ACKR3 (**Fig.4e**).

## Discussion

mRNA display has emerged as a powerful screening technology able to identify *de novo* ligands against GPCRs, such as the large extracellular domains of several class B receptors^45^. Here, we adapted RaPID to target a full-length class A GPCR target, showing the feasibility of this methodology to identify functional thioether-closed macrocyclic peptides against chemokine receptors in a nanodisc-based system. In Screen 1, nanodiscs were immobilized using biotinylated MSP leading to a mixture of empty- and CXCR4 nanodiscs during positive selection while direct capture of biotinylated CXCR4 in Screen 2 allowed for selection against only receptor nanodiscs. Both strategies generated peptides with distinct sequences and pharmacological properties. Even though the MSP-based approach had a lower positive over negative recovery than direct receptor capture, it showed the potential of RaPID selections against untagged GPCRs. Alternative selection strategies, including negative selection against related receptors, may be considered to improve target selectivity.

To determine the peptides’ ability to occupy the orthosteric pocket, we established two biophysical assays. While CXCL12 displacement broadly reflected the effects of most peptides on cell-based assays, neither peptide **H** nor **K** were shown to be significantly active in blocking *in vitro* mini-Gα_o_ recruitment. Surprisingly, our data showed that neither peptide **K** nor IT1t, a potent small-molecule inverse agonist for CXCR4, could outcompete CXCL12 from CXCR4 in our experimental setup. Since chemokines bind at the N-terminal region of a receptor through its chemokine recognition site 1 (CRS1) and through interactions deeper in the orthosteric pocket (CRS2)^46^, it is likely that **K** only competed for CRS2, as previously reported for IT1t^31^, thus allowing CXCL12 to remain bound at CRS1 without activating CXCR4. Importantly, this highlights how these two orthogonal biophysical approaches can complement each other to predict more accurately whether a receptor ligand will show cellular activity. Our results also suggest that macrocyclic peptides may be superior to small-molecule antagonists *in vivo*, as they may be more efficient at displacing increased local concentrations of chemokines resulting from receptor inhibition^47^.

The peptides also displayed diverse effects on chemokine receptor signaling. While **G** had preferential activity as a β-arrestin antagonist, **F** was consistently an antagonist in all assays. In addition, **H**, **I** and **K** all behaved as inverse agonists, just as IT1t, thereby reducing the constitutive G protein activation levels of CXCR4^33^. While all peptides binding CXCR4 were antagonists, seven of them showed agonism toward the related receptor ACKR3, similar to AMD3100^24^. Among the newly identified ACKR3 agonists, **B** was the most selective for ACKR3, **H** was a strong dual ACKR3 agonist and CXCR4 antagonist, and **K** was the most selective CXCR4 antagonist. The distinct activity of **G** and the different degrees of selectivity of **B**, **I** and **K** among related chemokine receptors further highlights the capacity of RaPID to generate ligands that either specifically modulate the complexity of the chemokine system or broadly target multiple related receptors.

Peptide **I** was the most potent CXCR4 antagonist across signaling assays. It also showed translational relevance, as it blocked CXCL12-driven T cell migration, and suppressed gp120-dependent viral fusion. Moreover, our selectivity screen showed that peptide **I** preferentially recognized CXCR4 among all 24 currently reported chemokine receptors. According to our results, treatment with peptide **I** at its IC_90_ (58 nM) should selectively antagonize CXCR4 without displaying off-target functions on any signaling chemokine receptor. Thus, peptide **I** is the most clinically relevant candidate derived from our two RaPID screens.

Using waterLOGSY, we identified that peptide **I** adopts a pose that leaves Trp9 and Leu6 exposed to the solvent, and Leu^10^ buried when interacting with CXCR4. These observations were consistent with the docking of peptide **I** with CXCR4. Our mutational analyses further provided experimental evidence for the interaction of peptides **I** and **H** with D187^ECL2^ in CXCR4 and E213^5.39^ in ACKR3, respectively. Mapping and quantifying the interaction sites of ACKR3 and CXCR4 with these peptides indicated how their modes of binding differed from those of CXCL12 and provided a molecular explanation for their experimental potencies.

A key aspect of drug discovery is the efficient and high-throughput screening of compounds to identify promising new drug candidates. Experimentally, RaPID selections can be achieved at a fast pace, as they require at most two weeks, provided that receptor purification and nanodisc reconstitution have been previously optimized. In addition, this method necessitates small amounts of purified receptor (< 100 µg), making it ideal for GPCRs whose expression and purification yields are low. Importantly, RaPID generates thioether-closed macrocyclic peptides, which overcome many limitations of other drug modalities^3^. The thioether bond in these macrocyclic peptides grants them higher rigidity, allowing for higher target selectivity, and higher resistance against proteolytic degradation compared to linear peptides^48^. The larger surface area in macrocyclic peptides also promotes stronger and more selective target binding compared to small-molecule compounds, thereby reducing off-target toxicity and enabling them to target traditionally undruggable proteins^49^.

Despite many years of drug discovery efforts and numerous clinical trials targeting chemokine receptors there are only five drugs approved to date. Failures have been attributed to heterologous post-translational modifications in the receptor’s N-terminal region affecting antibody binding, limited selectivity leading to off-target effects of small molecules, and the need to inhibit multiple receptors within the complex chemokine system^50^. We show that our macrocyclic peptides interact with the orthosteric pocket and ECLs and can be either receptor-selective or bind to multiple receptors. This suggests that macrocyclic peptides are a promising drug modality for the next generation of therapeutics, with promise to overcome the challenges of targeting the chemokine system.

Overall, our study establishes RaPID as a feasible and high-throughput strategy for discovering functional macrocyclic peptide ligands against chemokine receptors and other GPCRs using nanodisc systems. The identification of antagonists, inverse agonists and agonists with different selectivity profiles highlights the versatility of RaPID and supports its broader application to other GPCR targets that remain challenging for conventional ligand discovery.

## Methods

### Cell culture

Parental HEK293A cells (Thermo Fisher Scientific, Cat#R70507), HEK293T cells (Abcam), and Flp-In™-Jurkat cells (Thermo Fisher Scientific, Cat#R76207) were cultured at 37 °C in a humidified incubator with 5% CO₂. HEK293A cells were cultured using high-glucose Dulbecco’s Modified Eagle Medium (DMEM) containing GlutaMAX (Thermo Fisher Scientific, Cat#31966), supplemented with 10% fetal bovine serum (FBS) and 1% penicillin–streptomycin (Merck Life Science, Cat#P4333). HEK293A cell passage was routinely done by washing with phosphate-buffered saline (PBS) followed by treatment with 0.05% trypsin–EDTA (VWR, Cat#392-O459P). Flp-In™-Jurkat cells were cultured in RPMI 1640 medium (Thermo Fisher Scientific, Cat#A1049101), supplemented with 10% FBS and 1% penicillin–streptomycin. *Spodoptera frugiperda* (Sf9) insect cells (Expression Systems, Cat#94-001F) were cultured in suspension in ESF 921 Protein-Free Insect Cell Culture Medium (Clinisciences, Cat#96-001-01) at 27 °C with shaking at 140 rpm, and virus production was performed as previously described^18^.

### Cloning and receptor mutagenesis

The cDNAs encoding human CXCR4, ACKR3, and CXCR4 mutants fused to Rluc3 at their C-termini were cloned into the pcDNA3.1 expression vector. Constructs encoding N-terminally FLAG-SNAP-tagged CXCR4^51^, ACKR3^51^, and their respective mutants were cloned into the pcDNA5.1 vector. Mutations in CXCR4 and ACKR3 were introduced using the Stratagene QuikChange method with Phusion High-Fidelity PCR Master Mix with HF Buffer (Thermo Fisher, Cat#F531S) and *Dpn*l (Bionordika, Cat#R0176S). Venus-β-arrestin2, and GFP-mini-Gα_o_ constructs were generated using the FastCloning method^52^.

The Rluc3-tagged CXCR4 and ACKR3 constructs were kindly provided by Nikolaus Heveker (Université de Montréal, QC, Canada). Venus-mini-Gα_s/i_^53^, and the constructs used in heterotrimer dissociation assays (untagged Gα_i1_, β_1_, venus-γ_2_, and Nluc-tagged C-terminal tail of GRK3)^32^ were supplied by Nevin Lambert (Augusta University, GA, USA). Untagged mini-Gα_o_ was kindly provided by Arun Shukla. All constructs were verified by bidirectional DNA sequencing. The CD4, GAL4-VP16, and pGL4.31 plasmids were previously described^54^, and HIV gp120 was a gift from Joseph Sodroski (Dana-Farber Cancer Institute, Boston).

### Purification of CXCL12, MSP, BirA and GFP-mini-Gα_o_ from bacteria

Human CXCL12, MSP1D1, MSP1E3D1, BirA and GFP-mini-Gα_o_ were produced using previously established methods^19,55–57^. Briefly, the proteins were heterologously expressed in *Escherichia coli* BL21-Gold (DE3)pLysS using the pET-21 vector for CXCL12, the pET28a vector for MSP and BirA, or the pET-15b vector for GFP-mini-Gα_o_. The constructs contained an N-terminal His₈-(for CXCL12 and MSP) or His_6_-tag (for BirA and GFP-mini-Gα_o_). An enterokinase cleavage site was incorporated into the CXCL12 construct, whereas the remaining proteins contained a Tobacco Etch Virus (TEV) protease cleavage site.

Following expression, CXCL12 was recovered from inclusion bodies under unfolding conditions, while MSP, BirA and GFP-mini-Gα_o_ were extracted from soluble fractions of cell lysates. Initial purification of all proteins was performed using Ni-nitrilotriacetic acid (NTA) affinity chromatography. For CXCL12, MSP, and GFP-mini-Gα_o_, the affinity tag was subsequently cleaved using enterokinase or TEV protease, and the resulting proteins were subjected to a second Ni-NTA affinity binding step followed by reverse-phase chromatography. The purified proteins were then exchanged into their respected storage buffers: 20 mM acetate (pH 4.0) for CXCL12; 20 mM HEPES (pH 7.5), 150 mM NaCl and 5 mM sodium cholate for MSP; 20 mM Tris (pH 7.5), 100 mM NaCl for BirA; and 20 mM HEPES (pH 7.4), 500 mM NaCl, 10% Glycerol, 50 μM GDP and 1 mM MgCl_2_ for GFP-mini-Gα_o_. Finally, the protein preparations were concentrated by centrifugal filtration and stored at −80 °C until further use.

For Screen 1, MSP1E3D1 was biotinylated by incubation with a four-fold molar excess of EZ-Link NHS-PEG4-Biotin (ThermoFisher Scientific, Cat#A39259) for 1 h at room temperature. The reaction was then quenched with 5 mM of TRIS (pH 7.5).

### Expression and purification of CXCR4

Following previously established protocols^58^, we expressed wildtype human CXCR4 (residues 2–352) in Sf9 insect cells using the Bac-to-Bac baculovirus expression system (Thermo Fisher Scientific). The cDNA of CXCR4 was cloned into the pFastBac1 vector, containing an N-terminal polyhedrin (PolH) promoter, a hemagglutinin (HA) signal peptide, and three C-terminal affinity tags: His₁₀, FLAG and AviTag™. Cells expressing CXCR4 were harvested 48 h post-transduction, and membrane fractions were isolated using by Dounce homogenization in a hypotonic buffer. The CXCR4-containing membranes were subsequently subjected to repeated high-salt washes. EDTA-free protease inhibitor cocktail (Sigma-Aldrich, Cat#5056489001) was included during CXCR4 purification.

Before detergent-mediated extraction, the membranes were incubated with 100 µM IT1t dihydrochloride (MedChemExpress, Cat#HY-101458A). Membrane proteins were solubilized for 1 h in a buffer containing 50 mM HEPES (pH 7.5), 400 mM NaCl, and 0.75% (w/v) *n*-dodecyl-β-D-maltopyranoside (DDM; Anatrace, Cat#D310), without addition of cholesteryl hemisuccinate (CHS). Following solubilization, the insoluble fraction was removed by centrifugation at 50,000 × *g* for 30 min. The resulting supernatant was adjusted to 20 mM imidazole, and incubated overnight at 4 °C with TALON® Superflow™ resin (Cytiva, Cat#28-9575-02) for immobilized metal affinity purification. DNase I (Thermo Scientific, Cat#90083) was added to a final activity of 1 U mL⁻¹ during the binding step.

After overnight incubation, the resin was washed using detergent-containing buffers with successively decreasing DDM concentrations. Bound CXCR4 was eluted with 500 mM imidazole, and the eluate was desalted using PD MiniTrap G-25 columns (Cytiva, Cat#28918007). The exchanged buffer contained 50 mM HEPES (pH 7.5), 400 mM NaCl, and 0.025% (w/v) DDM. The purified protein was concentrated using 100-kDa molecular weight cut-off protein concentrators (Thermo Fisher Scientific, 88533), and its purity was assessed by SDS-PAGE. Protein concentration was measured spectrophotometrically at 280 nm using the extinction coefficient of 65,820 M^−1^ cm^−1^. Finally, the purified protein was snap-frozen in liquid nitrogen and stored at −80 °C until use.

### Nanodisc reconstitution and CXCR4 biotinylation

Nanodiscs were prepared using a method based on^18^. 1-palmitoyl-2-oleoyl-*sn*-glycero-3-phosphocholine (POPC; Merck Life Science, Cat#850457C) and 3β-hydroxy-5-cholestene (cholesterol; Merck Life Science, Cat#C3045) were first dissolved in chloroform and mixed at a molar ratio of 9:1 (POPC:cholesterol). The solvent was evaporated under a gentle nitrogen stream, and the resulting lipid film was resuspended in cholate-containing buffer to achieve a final lipid concentration of 100 mM. The buffer consisted of 50 mM HEPES (pH 7.5), 150 mM NaCl, and 24 mM sodium cholate.

To promote the incorporation of individual receptor molecules into each nanodisc, two membrane scaffold protein (MSP) variants, MSP1D1 and MSP1E3D1, were employed, yielding particles with approximate diameters of 10 or 12 nm, respectively^59^. For Screen 1, nanodiscs contained purified N-terminally FLAG-tagged CXCR4, biotinylated MSP1E3D1 scaffold protein, and lipids at a molar ratio of 0.1:1:105 (CXCR4:MSP1E3D1:lipid). For Screen 2, nanodiscs were reconstituted by mixing purified CXCR4, non-biotinylated MSP1D1 scaffold protein, and lipids at a molar ratio of 0.1:1:76 (CXCR4:MSP1D1:lipid). In both cases, a final cholate concentration above 20 mM was maintained. Empty nanodiscs were reconstituted as described for Screen 1, where the receptor solution was replaced by nanodisc buffer (25 mM HEPES, pH 7.5, 50 mM NaCl).

The assembly mixtures were incubated at room temperature for 20–30 min with gentle agitation. Detergent removal was then initiated by adding Bio-Beads SM-2 resin (Bio-Rad, Cat#152-3920), followed by overnight incubation at 4 °C to facilitate spontaneous nanodisc formation. The resin was removed after nanodisc reconstitution. For Screen 2, the resulting CXCR4-containing nanodiscs were then enzymatically biotinylated using BirA. The reaction was performed at room temperature for 2 h without agination, using 0.0125 mg/mL BirA, 0.2 mM biotin, 10 mM ATP and 10 mM MgCl_2_.

Nanodisc samples were purified by size-exclusion chromatography on a ProteoSEC Dynamic 11/30 6-600 HR column (Protein Ark/Bionordica, Cat#SEC-D-11/30-6-600) equilibrated with nanodisc buffer. Fractions containing nanodiscs were combined and assessed by SDS-PAGE. The purified nanodisc preparations were concentrated to final concentrations of approximately 1–30 μM MSP, and stored at 4 °C for short-term use, or at −80 °C for long-term storage.

### RaPID

tRNA synthesis, mRNA library synthesis, flexizyme-based genetic code reprogramming and RaPID selections were performed as described in^60^, with the following modifications. The initial mRNA library consisted of a (NNK)_n_ degenerate codon library with n = 4–15, in ratios 0.001:0.01:0.01:0.1:1:1:1:1:1:1:1:1. As nanodiscs are employed during selection, Tween-20 was reduced to a concentration of 0.01% (instead of the typical 0.05%) in the blocking and selection buffers. The blocking buffer contained 50 mM Hepes-KOH (pH 7.6), 100 mM NaCl, 1 mM DTT, 0.01% Tween-20, 0.1% UltraPure BSA. The selection buffer had the same composition without UltraPure BSA. Assuming 100% protein purity, loading capacity was estimated to be 0.5 or 1.0 pmol/µl for Streptavidin Dynabeads M280 beads (ThermoFisher) for Screens 1 & 2, respectively. 200 nM protein target was used for all RaPID screens, and incubation and washing of the libraries was conducted on ice.

### Fluorescence polarization-based CXCL12 competition assay

Direct CXCL12 binding to CXCR4 in DDM was evaluated using C-terminally AlexaFluor488-labeled CXCL12 (Protein Foundry, Cat#PFP001T-AZ488) in a fluorescence polarization assay. For this, CXCL12-488 was initially dissolved in double-distilled water (ddH_2_O) at a stock concentration of 2 µM. CXCL12-488 was then diluted to a concentration of 25 nM in reaction buffer (50 mM Hepes, pH7.4; 400 mM NaCl; 0.025% DDM; 10% glycerol; 0.1% fatty acid-free BSA). CXCR4 was diluted serially two-fold in reaction buffer. 25 µl of each step in the CXCR4 dilution series (at two times its final concentration) was added onto flat-bottomed 96-well microplate with reduced non-specific binding (Greiner, Cat#655900). Next, 25 µl CXCL12 was added to the wells for a final concentration of 12.5 nM, and fluorescence polarization was measured using a PHERAstar microplate reader (BMG Labtech).

We evaluated CXCL12 displacement as an effect of peptide competition for CXCR4 binding. For this, a fixed final concentration of 500 nM CXCR4 was prepared in reaction buffer and added to the wells in 25 µl. Next, 12.5 µl peptides or IT1t was added for a final concentration of 5 µM and allowed to incubate with CXCR4 for 10 min. 12.5 µl CXCL12 (final concentration = 12.5 nM) was then added to the wells, incubated for 10 min, and fluorescence polarization was measured using a PHERAstar microplate reader.

### *In vitro* mini-Gα_o_ binding to CXCR4

A fluorescence saturation binding assay (FSBA) was used to detect *in vitro* G protein binding. This method was performed as described in^29^, and was originally based on^61^. His_10_-tagged CXCR4 was reconstituted into nanodiscs containing untagged MSP1D1 and POPC:cholesterol (9:1) at a molar ratio of 0.1:1:76 (CXCR4:MSP1D1:lipid). To assess non-specific binding, empty nanodiscs were prepared using His_8_-tagged MSP1D1 and the same lipid composition.

For each experimental condition, 10 µl of TALON® Superflow™ resin were transferred to a new tube and centrifuged at 100 × g for 5 min. The resin was washed twice with a binding buffer consisting of 25 mM HEPES (pH 7.5), 50 mM NaCl, and 0.1% BSA, (w/v). Following removal of the supernatant, 50 pmol of CXCR4-containing nanodiscs was added to each tube. The resin was then washed twice with binding buffer.

GFP-mini-Gα_o_ was then added at a final concentration of 5 µM to the tubes containing the resin. For each sample, one of the following was added: nanodisc buffer, 0.5 µM CXCL12, 5 µM peptide, or 5 µM IT1t. The mixtures were incubated for at least 30 min at room temperature. Following incubation, the samples were maintained at 4 °C to reduce rapid dissociation. The resin was then pelleted, and the supernatant was removed using a multi-channel pipette before addition of fresh binding buffer. This wash was repeated two more times.

To elute CXCR4 nanodiscs from the resin, the beads were incubated in binding buffer supplemented with 300 mM imidazole for at least 15 min. The resulting supernatants were then collected, and fluorescence emission was measured at 509 nm using a PHERAstar microplate reader (BMG Labtech).

### Cell viability

Following the protocol in “BRET-based signaling assays”, each peptide was incubated at a concentration of 5 µM in cell culture buffer for 24 h. After this, the supernatant was removed, the cells were washed twice with PBS, and cell viability was tracked using CellTiter-Glo® Luminescent Cell Viability Assay (Promega, Cat#G7570).

### BRET-based signaling assays

For bioluminescence resonance energy transfer (BRET) assays, HEK293A cells were plated at a density of 30,000 cells per well in while, poly-D-lysine-coated 96-well plates (Greiner. Cat#655083) and co-transfected with the amounts of DNA constructs indicated below using polyethyleneimine (PEI) at a DNA:PEI ratio of 1:2 (w/w). After 48 hours, the culture medium was removed, and the wells were rinsed twice with PBS. BRET buffer (PBS supplemented with 5mM glucose) was subsequently added to each well at a volume of 80 µl to test agonism, or 70 µl to test antagonism.

Coelenterazine H (Nanolight technology, Cat#301–10) was added at 10 µl per well for a final concentration of 5 μM, followed by an 8-min incubation. For antagonist assays, 10 µl of the peptide or small-molecule compound was added at the indicated concentrations and incubated simultaneously with the substrate. Following substrate (and antagonist) incubation, the cells were stimulated with agonist: 10 µl peptides or small-molecule compounds at the indicated concentrations to test agonism, or 10 µl CXCL12 (for a final concentration of 10 nM) to test antagonism. The plate was then read at 37 °C using an EnVision Plate Reader (Revvity), and BRET ratios were calculated from the measured signals:

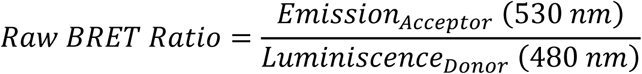

Concentration-response data was analyzed in GraphPad Prism (version 11.0.2) using four-parameter nonlinear regression with a variable slope. The upper and lower asymptotes of the fitted curves were used to determine efficacy and baseline responses, respectively. Compound potency was derived from the fitted Log(EC_50_) or Log(IC_50_) values of the fits.

The DNA constructs, as well as their transfected amounts, are described below for the individual signaling assays.

(I) Cell-based mini-Gα_s/i_ coupling assay^53^. HEK293A cells were co-transfected with 2.5 ng/well C-terminally Rluc3-tagged CXCR4 and 10 ng/well N-terminally venus-tagged mini-Gα_s/i_, where empty pcDNA3.1 as added for a final DNA amount of 50 ng per well of a 96-well plate.
(II) Heterotrimer dissociation-based G protein activation assay^32^. HEK293A cells were co-transfected with 5 ng/well N-terminally SNAP-tagged receptor, 36 ng/well untagged Gα_i1_, 12.5 ng/well Gβ_1_, 12.5 ng/well venus-Gγ2, and 2.5 ng/well Nluc-tagged C-terminal tail of GRK3. The final amount of DNA in this assay was 68.5 ng/well.
(III) Direct β-arrestin2 recruitment assay^51^. HEK293A cells were co-transfected with 2.5 ng/well of C-terminally Rluc3-tagged receptor and 10 ng/well venus-β-Arrestin2, where empty pcDNA3.1 vector was added for a total transfected DNA amount of 50 ng/well.

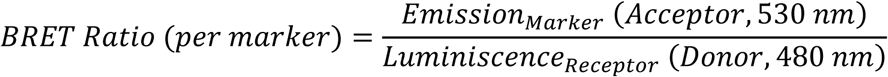

All datasets were analyzed statistically using ordinary one- or two-way ANOVA followed by Dunnett’s multiple-comparisons test.

### Receptor internalization

Detection of receptor internalization was performed using diffusion-enhanced resonance energy transfer (DERET) using SNAP-based receptor labeling. Energy transfer was measured between a terbium fluorophore probe SNAP-Lumi4-Tb (Revvity, Cat#SSNPTBD) and the acceptor fluorescein-O-acetic acid (Sigma-Aldrich, Cat#88596) acceptor in the extracellular milieu^62^. HEK293A cells were seeded at a density of 20,000 cells per well into poly-D-lysine-coated white 384-well plates (In Vitro, Cat#GR-781201) and co-transfected with 10 ng of N-terminally FLAG-SNAP-tagged GPCR plasmid together with 30 ng of pcDNA3.1, for a total of 40 ng DNA per well. Transfections were performed using Lipofectamine™ 2000 (Thermo Fisher Scientific, Cat#11668027) at a DNA-to-reagent ratio of 1:2 (w/v). 24 h later, culture medium was replaced with Opti-MEM™ I Reduced Serum Medium containing 100 nM SNAP-Lumi4-Tb, where cells incubated for 1 hour at 4 °C. Following labeling, cells were washed six times with internalization buffer, composed of Hanks’ Balanced Salt Solution (HBSS), 1 mM CaCl₂, 1 mM MgCl₂, 20 mM HEPES, and 0.1% (v/v) bovine serum albumin (BSA) at pH 7.4. For internalization, fluorescein-O′-acetic acid (Merck Life Science, Cat#88596) was added in 10 µl to the wells at a final concentration of 25 μM, followed by 10 μl buffer or ligand. The assay was read immediately at 37 °C using an EnVision Plate Reader (Revvity). The following formula was used to calculate internalization ratios:

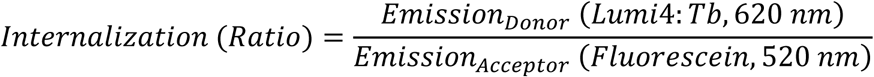

Internalization was quantified using the area under the curve in GraphPad Prism (v11.0.2). Statistical differences between groups were evaluated using one-way ANOVA followed by Dunnett’s multiple-comparison post hoc test.

### Trans-well cell migration

Wild-type Flp-In™-Jurkat cells were cultured to reach a density of 2.0 × 10^6^ cells/mL. A sample with enough cells for each experiment was taken, centrifuged to pellet the cells at 200 × g for 5 min, after which the supernatant was removed. The cells were then resuspended in 5 mL PBS, centrifuged at the same speed, and the supernatant was removed. The cells were then resuspended in migration buffer (RPMI 1640 medium supplemented with 2% FBS and 1% penicillin–streptomycin), and diluted to 1.0 × 10^6^ cells/mL. Wells in a 24-well plate (lower chamber) were filled with 600 µl containing migration buffer with or without 10 nM CXCL12, 5 µM peptides or IT1t. Corning® 6.5-mm Transwell® inserts with 5.0-µm Pore Polycarbonate Membranes (Corning, Cat#3421) were carefully placed over the lower chamber, avoiding bubble formation. 100 µl of the cell suspension was added on top of the inserts (upper chamber) and were allowed to migrate for 3 h. 20 min before harvest, 60 µl EDTA was added to the lower chamber for a final concentration of 5 mM to detach cells loosely adhered to the membrane. Cells were harvested from the lower chamber, centrifuged at 200 × g for 5 min, and 560 µl supernatant was removed. Cells were then resuspended in the remaining 100 µl supernatant, and 25 µl were added to a flat-bottomed black plate. 25 µl CellTiter-Glo® (Promega, Cat#G7570) solution was then added to all wells, cells were allowed to lyse for 10 min in the dark with shaking at 300 rpm. Cell viability was then measured using an EnVision Plate Reader (Revvity).

### HIV gp120-mediated cell-cell fusion assay

The cell-cell fusion assay detected the fusion of HIV gp120-expressing effector cells with CD4/CXCR4-expressing target cells, as described in^54^. For the effector cells, 800,000 HEK293 cells/well were seeded in 6-well plates. For the target cells, 20,000 HEK293 cells/well were seeded in poly-D-lysine-coated white 96-well plates (Greiner bio-one). Cells were incubated overnight and transfected the following day using Lipofectamine™ 2000. The effector cells were co-transfected with 0.5 µg HIV gp120 and 1.1 µg GAL4-VP16 per well. Target cells were co-transfected with 30 ng CD4, 5 ng SNAP-CXCR4, and 40 ng pGL4.31 per well. The day after transfection, effector cells were detached using EDTA, resuspended in antibiotic-free culture medium, and mixed with ligand or vehicle at the indicated concentrations. Culture medium was then removed from the target cells and replaced with 100 µL culture medium containing 200,000 effector cells in the respective ligand; cells were co-cultured overnight. After co-culture, cells were washed with 100 µL HBSS++ (Gibco) and pGL4.31-derived luciferase activity was detected using the Firefly Assay Buffer from^63^, in an EnVision Plate Reader (Revvity).

### Chemokine receptor selectivity screening

Ligand-induced β-arrestin recruitment to classical and atypical chemokine receptors was monitored by Nanoluciferase complementation assay (NanoBiT). In brief, 6 × 10^6^ HEK293T cells were plated in 10-cm culture dishes and 24 hours later transfected with vectors encoding receptors C-terminally tagged with SmBiT and human β-arrestin-1 N-terminally tagged with LgBiT. The only exception was CXCR4, which was tagged C-terminally with LgBiT and used with human β-arrestin-1 N-terminally tagged with SmBiT. 24 hours after transfection, cells were harvested, incubated 25 minutes at 37 °C with coelenterazine H (4 ng/ml) and distributed into white 96-well plates (1 × 10^5^ cells per well). Peptide **I** (5 μM) was then added, and luminescence was measured with a Mithras LB940 plate reader. Next, peptide antagonist activity was determined by measuring luminescence after addition of a positive control chemokine at a concentration corresponding to the EC_50_ value for each receptor (**Table S3**).

The results were expressed as percentage of signal obtained with a known endogenous agonist chemokine (200 nM) listed in the IUPHAR repository of chemokine receptor ligands (positive control chemokine) for agonist mode, or as percentage of signal obtained with the positive control chemokine at a concentration corresponding to the EC_50_ value for each receptor for the antagonist mode.

### NanoBRET binding competition assay

The binding of the different ligands to CXCR4, ACKR3 or ACKR5 was monitored by NanoBRET. HEK293T cells were transfected with vectors encoding the receptor N-terminally tagged with Nanoluciferase. 24 hours after transfection, cells were distributed into white 96-well plates (0.8 × 10^5^ cells per well) and incubated for 5 minutes on ice. Subsequently, unlabeled peptide at the indicated concentrations and AZDye568-labeled CXCL12 (2 nM (ACKR3), 30 nM (ACKR5) and 10 nM (CXCR4)) were added. After 2-hour incubation on ice, the Nanoluciferase substrate coelenterazine H (4 ng/ml) was added and donor (450/80 BP) and acceptor (610 LP) emissions were measured on PHERAstar plate reader. Results were expressed as BRET ratio (acceptor/donor emission). Cells incubated with CXCL12-AZ568 alone were used to define 100% BRET binding.

### NMR-based ligand interaction analysis

WaterLOGSY experiments were performed using CXCR4 POPC MSP1D1 nanodiscs; reconstitution was performed as described above, omitting cholesterol. The waterLOGSY sample contained 250 µM peptide **I** and 2.5 µM CXCR4 nanodiscs in NMR nanodisc buffer (nanodisc buffer with 10% D_2_O (v/v)). The two control samples contained 250 µM peptide **I** in NMR nanodisc buffer and 250 µM peptide **I** with 2.5 µM empty POPC MSP1D1 nanodiscs in NMR nanodisc buffer respectively.

^1^H and waterLOGSY spectra were collected of all samples. All spectra were collected in standard 5mm NMR tubes on a 750 MHz Bruker Avance III HD spectrometer equipped with a 5mm TCI cryoprobe H-C/N-D optimised for ^13^C. waterLOGSY experiments were acquired with the ephogsygpno.2 pulse program from the standard Bruker library. A mixing time of 1.5 seconds was used and 1024 scans collected for an experiment time of approximately 1 hour per condition. Spectra were processed in TopSpin 5.0 and visualised using bespoke python scripts. WaterLOGSY were phased such that negative NOEs were phased positively as per convention^64^. Due to the limited NOE enhancement observed in the buffer only control, comparisons were primarily done using the empty nanodisc control.

Spectral assignment of peptide **I** was performed using peptide dissolved in water (10% D_2_O) to better resolve the backbone amides and eliminate interference from buffer peaks. TOCSY, ROESY and HSQC spectra were collected for assignment. Spectra were processed in TopSpin 5.0 (Bruker) and assigned in CARA (cara.nmr.ch)^65^. In some cases, multiple states of pepide **I** were observed for residues. In such instances, only one state was assigned. Assignments were transferred by comparison to peptide in buffer. Assignment table is provided in **Table S4**.

### Structural co-folding and receptor-ligand contact analysis

Complexes of human CXCR4 and ACKR3 with CXCL12, peptide H, or peptide I were predicted using AlphaFold 3^40,66^ and subsequently relaxed using Rosetta FastRelax^42^ (Rosetta version 2024.09). One relaxed structure per receptor–ligand combination was analysed, yielding six models. Receptors and ligands occupied chains A and B, respectively. Peptides **H** and **I** were represented as non-standard ligand residues, whereas CXCL12 was represented as a polypeptide chain.

Receptor–ligand interactions were identified using PLIP version 3.0.0^43^ with default interaction-specific geometric criteria. CXCL12 was analysed in peptide mode, specifying chain B, whereas H and I were analysed using ligand detection. Interactions between receptor and ligand chains were extracted from the XML reports and classified as hydrogen bonds, salt bridges, cation–π interactions, π-stacking interactions, or hydrophobic contacts. For interaction matrices and composition counts, each receptor-residue/interaction-type combination was counted once. Residues participating in multiple interaction types contributed to each corresponding category.

Receptor residues were annotated using GPCRdb-derived generic numbering^67^; extracellular loop residues without a specific generic annotation were labelled by loop and receptor residue number. N-terminal contacts were excluded. For each receptor, H and I were compared separately with CXCL12, and contacted positions were classified as shared, peptide-only, or CXCL12-only. Each position was counted once in these comparisons, irrespective of interaction type or multiplicity, without common-core subtraction. Contact tables and plots were generated using Python, pandas, and Matplotlib. Structures were visualized using PyMOL^68^ .

## Data analysis

Dose–response curves were analyzed using a four-parameter nonlinear regression model of log-transformed agonist concentrations versus response, incorporating a variable slope. Variability between experimental days was minimized by comparing data from independent experiments using ordinary one-way ANOVA, followed by Dunnett’s multiple-comparison test. All statistical analyses were performed in GraphPad Prism version 11.0.2.

## Data availability

All data are available upon request to.

## Acknowledgements

This work was supported by Villum Fonden grant 00025326 (M.G.), Independent Research Fund Denmark grant 3103-00230B (M.G.), Danish Cancer Society grant R389-A22939 (F.S.-P.), Novo Nordisk Foundation grants NNF22OC0080625 (M.M.R.), NNF24OC0094987 (M.M.R.), NNF25OC0100883 (J.M.R.) and NNF19OC0054441 (J.M.R.), European Research Council (ERC) under Horizon Europe grant agreement no. 101055152 (M.M.R.), the Carlsberg Foundation grant CF20-0248 (A.S.H.), the European Commission grant agreement no. 101052342 (A.S.H) and by the Luxembourg Institute of Health (LIH) through the NanoLux Platform, the Cancer Foundation Luxembourg, Luxembourg National Research Fund (INTER/FNRS CXCL12 20/15084569, CORE IMPACTT C23/BM/18068832, ACKROS C25/BM/19563724, INTER/AUDACE 25/19352183). The authors would like to thank Manuel Counson and Nadia Beaupain for technical assistance. NMR data were recorded at cOpenNMR, an infrastructure facility funded by the Novo Nordisk Foundation (NNF18OC0032996)

## Contributions

F.S.-P., J.M.R., and M.G. were responsible for study conceptualization. F.S.-P, M.Matic., F.F., M.M.R., M.S., A.C., A.S.H., J.M.R., and M.G. were responsible for experimental design. F.S.-P., P.B.R., F.F., M.Meyrath., and R.R. conducted wet-lab experiments, data analysis, and visualization. M.Matic. conducted dry-lab experiments. F.S.-P wrote the original draft, which was revised and polished by M.M.R., M.S., A.C., A.S.H., J.M.R., and M.G.

