## Extended Data Figures S1-S5 and Table S1-S4 for "*De novo* macrocyclic peptide discovery produces potent class A GPCR ligands with diverse pharmacology"

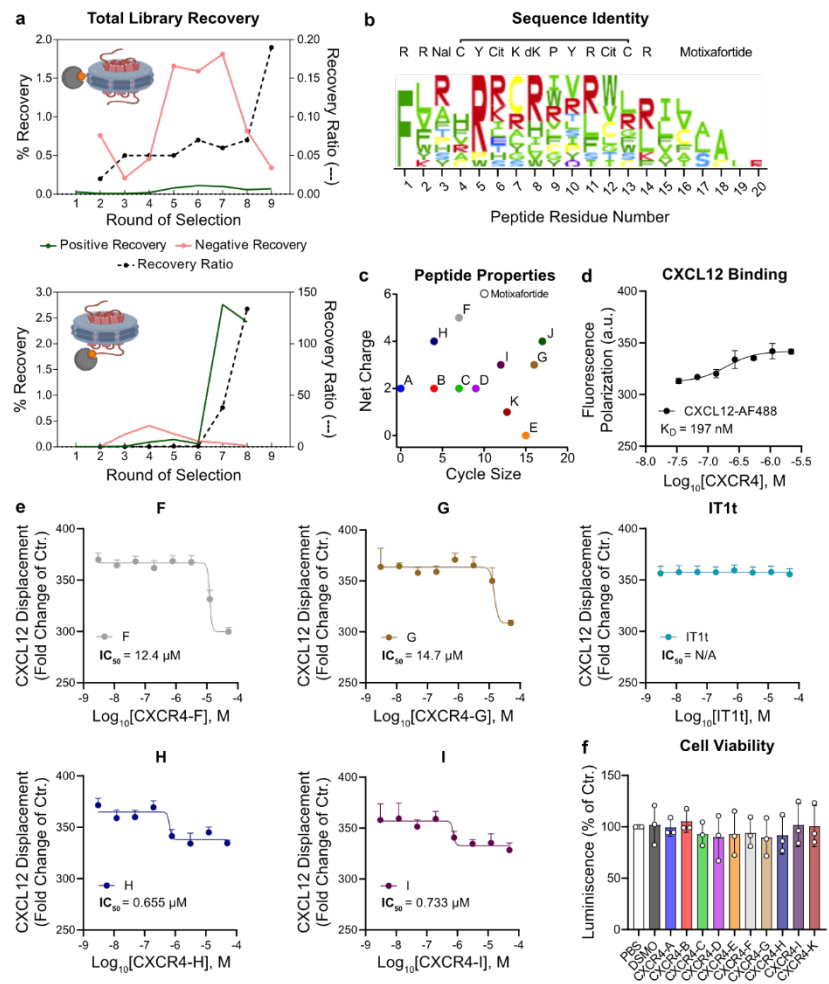

**Fig. S1: Discovery of novel peptides targeting CXCR4 using RaPID.** **a**, Bulk DNA-tagged library recovery per RaPID round measured by real-time quantitative PCR (RT-qPCR) for both screens. Positive and negative recovery refers to the amount of DNA present in CXCR4-, or empty nanodisc-coated beads, respectively. **b**, Sequence alignment of all eleven unique peptides selected in both screens compared to motixaforide, depicting its cyclization site between the Cys4 and Cys13. **c**, Comparison of the diverse charge and cycle size obtained from both RaPID screens compared to motixafortide. **d,e**, Dose-response curves of CXCL12-AF488 binding to CXCR4 (**d**), or its displacement by the identified peptides (**e**). **f**, Effects of 24-h peptide treatment (5 μM) on HEK293A cell viability, detected using CellTiter-Glo® Luminescent Cell Viability Assay. (**f**) No significant differences between PBS and the remaining groups, determined by one-way ANOVA. (**d-f**) N = 3 independent experiments, performed in technical triplicates. All data are shown as mean ± SD.

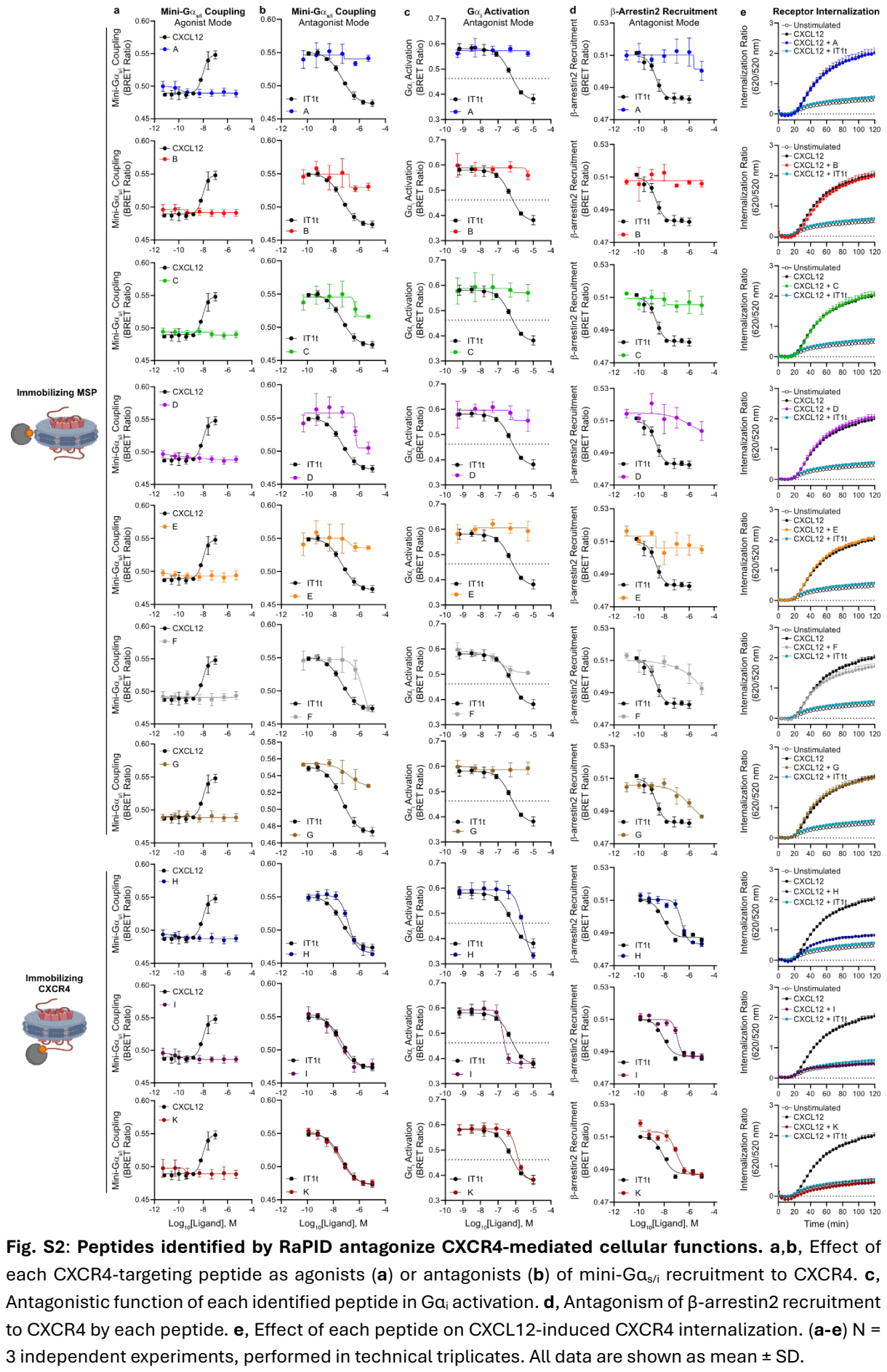

Fig. S2: Peptides identified by RaPID antagonize CXCR4-mediated cellular functions. a,b, Effect of each CXCR4-targeting peptide as agonists (a) or antagonists (b) of mini-Gαs/i recruitment to CXCR4. c, Antagonistic function of each identified peptide in Gαi activation. d, Antagonism of β-arrestin2 recruitment to CXCR4 by each peptide. e, Effect of each peptide on CXCL12-induced CXCR4 internalization. (a-e) N = 3 independent experiments, performed in technical triplicates. All data are shown as mean ± SD.

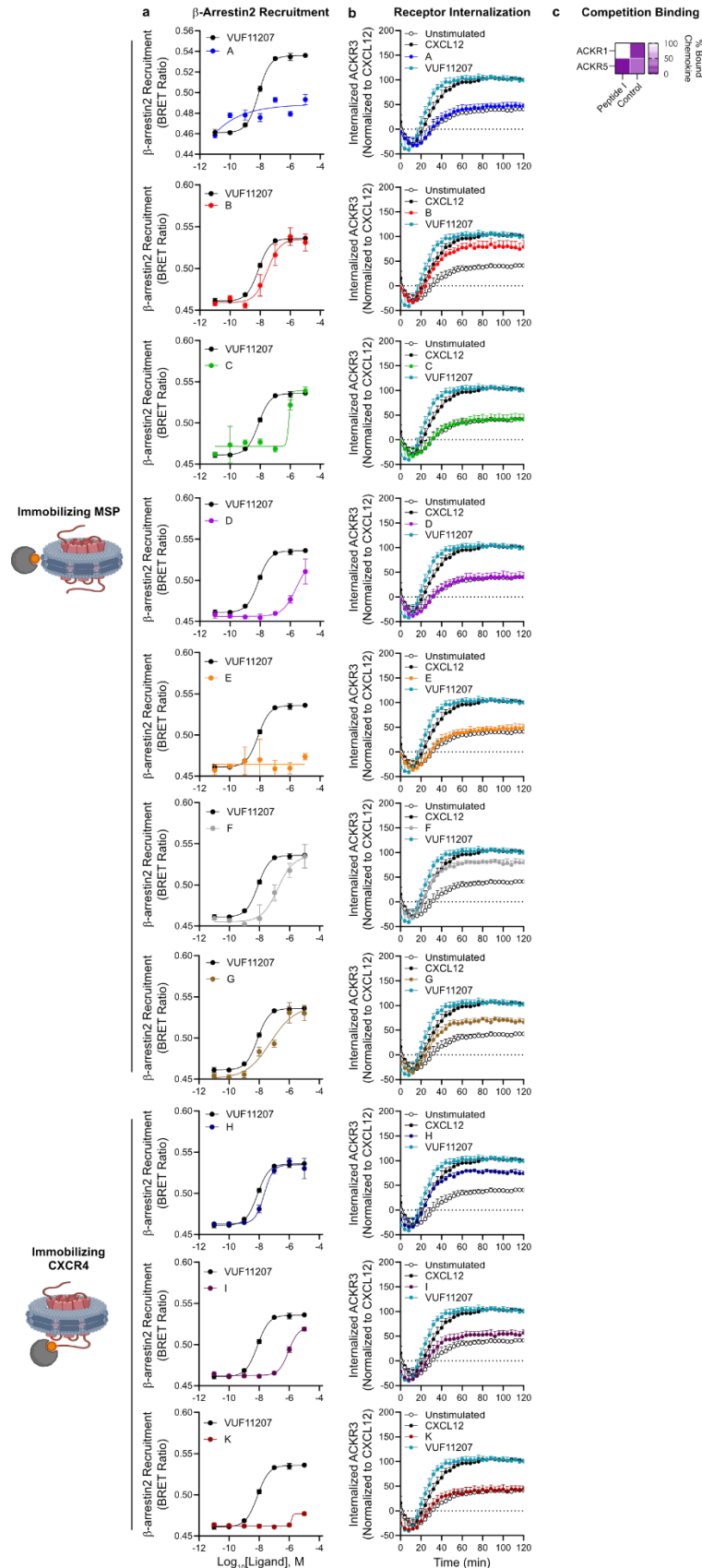

**Fig. S3: Novel CXCR4 antagonists show differential selectivity towards CXCR4.** **a,b,** Effect of each CXCR4-targeting peptide as potential ACKR3 agonists in β-arrestin2 recruitment to ACKR3 (**a**), or receptor internalization (**b**). **c,** Displacement of labeled chemokine by peptide **I** from N-terminally Nluc-tagged ACKR1 or ACKR5. N = 3 independent experiments, performed in technical triplicates. All data are shown as mean ± SD.

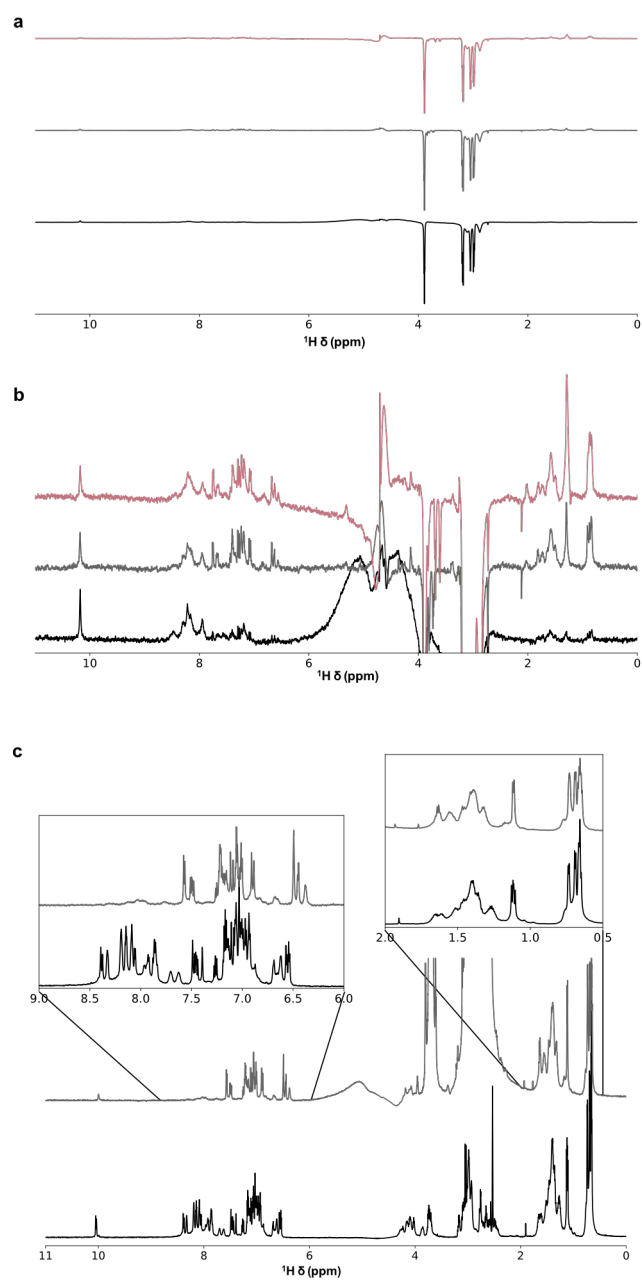

**Fig. S4: NMR spectra of peptide I.** **a,b**, Full waterLOGSY spectra of peptide I in buffer (black), with empty POPC nanodiscs (grey), and CXCR4 containing POPC nanodiscs (red). **(a)** Zoomed-out view showing all peaks; **(b)** zoomed-in showing all peptide peaks. **c**, Comparison of $^1\text{H}$ spectra of peptide I in buffer (grey) and water (black). Zoom-ins highlight the amide and aromatic, and the aliphatic spectral regions. The aliphatic region remains largely unchanged between the two solvents. However, in addition to the sharper amides in water, some chemical shift changes can be observed in water.

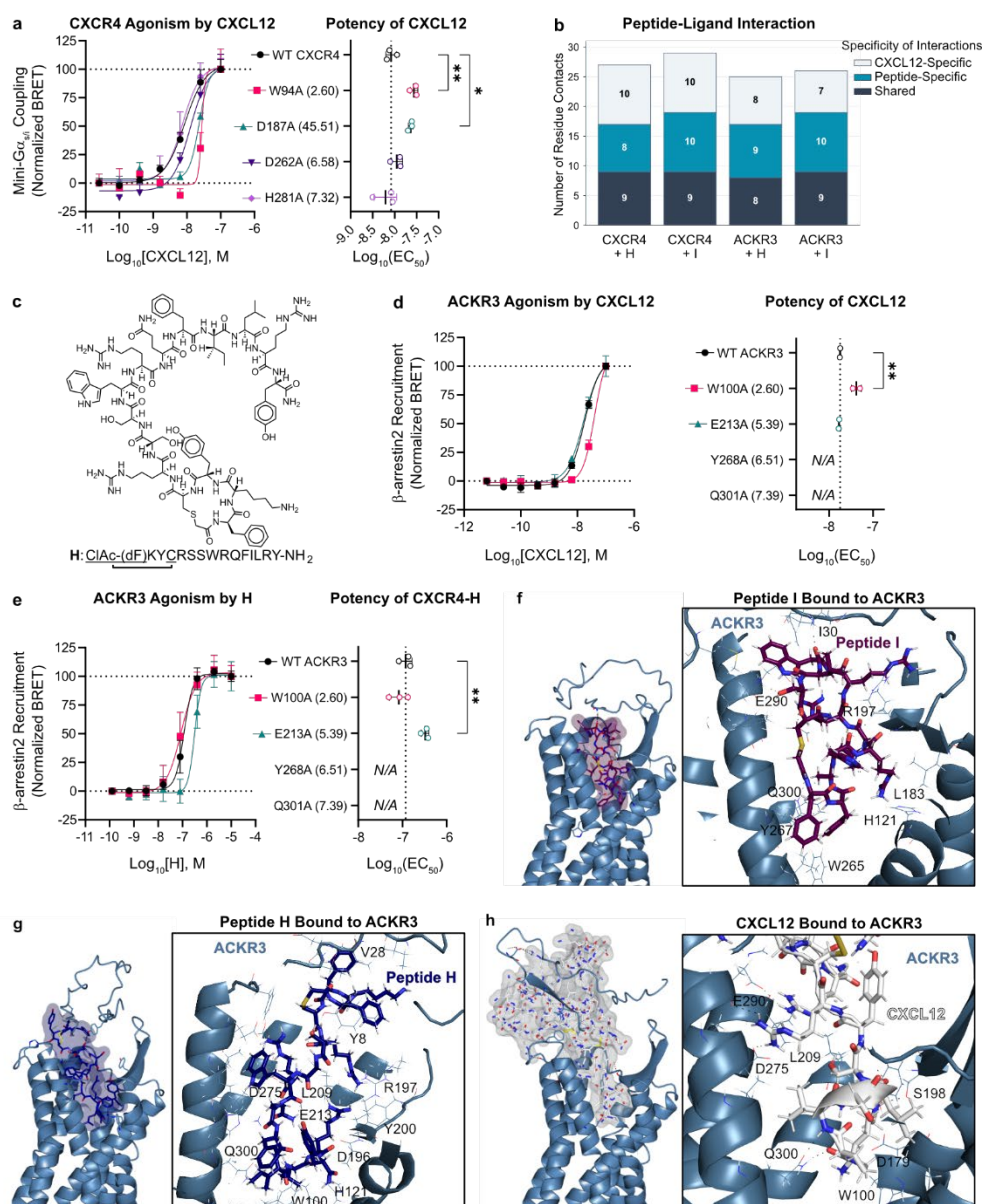

**Fig. S5: Most potent CXCR4 inverse agonists specifically interact in the orthosteric pocket.** **a**, BRET-based mini-G$\alpha_{s/i}$ coupling assay detecting direct recruitment to CXCR4 mutants in response to CXCL12. The results are shown as dose-response curves (left), and the quantification of CXCL12 potency for each mutant. **b**, Quantification of specific interactions between CXCR4 and peptide **I**, or ACKR3 and peptide **H**, considering those involved in CXCL12 binding. Total receptor-CXCL12 interactions (a total of 19 for CXCR4, and 16 for ACKR3, respectively) were separated into two groups: those shared with each peptide, and those that only account for CXCL12 binding. **c**, Structure and sequence of peptide **H**, depicting the cyclization site between the N-terminal CIAC-(dF) and Cys4. **d**, Detection of CXCL12-induced β-arrestin2 recruitment to ACKR3 mutants, shown for each as dose-response curves (left), and the quantification of CXCL12 potency. **e**, Effect of peptide **H** on the induction of β-arrestin2 recruitment to ACKR3 mutants. For each mutant, dose-response curves (left) and peptide **H** potency are shown. **f,g,h**, Predicted mode of binding between ACKR3 (in light blue) and peptide **I** (**f**, in purple), peptide **H** (**g**, in dark blue), or CXCL12 (**h**, in gray). (Left) View of the extracellular site of ACKR3 (in ribbon representation), and the van der Waals radius of each ligand (in surface representation). (Right) Residues at a maximum distance of 2.5 Å from each ligand, including hydrogen bonds. Images created with PyMOL. (**a,c,e**) The asterisk symbols indicate statistically significant differences (\*$p < 0.05$, \*\*$p < 0.01$) between the indicated groups, determined by one-way ANOVA. Only significant differences to the control group are shown. $N = 3$ independent experiments, performed in technical triplicates. All data are shown as mean ± SD.

**Table S1: List of RaPID-selected peptides compared to motixafortide.**

| Ligand | Sequence* | Size | Cycle | Net Charge |
| --- | --- | --- | --- | --- |
| **A** | ClAc-(dF)APKTLYNHGQLGPR-NH<sub>2</sub> | 15 | 0 | +2 |
| **B** | ClAc-(dF)ALCQSYVLRQLVR-NH<sub>2</sub> | 14 | 4 | +2 |
| **C** | ClAc-(dF)ARNWLQVWQLGLR-NH<sub>2</sub> | 14 | 7 | +2 |
| **D** | ClAc-(dF)ALILRVTCERGTQLVR-NH<sub>2</sub> | 17 | 9 | +2 |
| **E** | ClAc-(dF)ELTKWNANLPTGIC-NH<sub>2</sub> | 15 | 15 | 0 |
| **F** | ClAc-(dF)WRFRRCLGLRWWRIA-NH<sub>2</sub> | 16 | 7 | +5 |
| **G** | ClAc-(dF)LHRWTYHYPRTSRAC-NH<sub>2</sub> | 16 | 16 | +3 |
| **H** | ClAc-(dF)KYCRSSWRQFILRY-NH<sub>2</sub> | 16 | 4 | +4 |
| **I** | ClAc-(dF)YAHRLRRWLSC-NH<sub>2</sub> | 12 | 12 | +3 |
| **J** | ClAc-(dF)VPRRARFTLGLKFVSC-NH<sub>2</sub> | 17 | 17 | +4 |
| **K** | ClAc-(dF)VSARHGHYIWQC-NH<sub>2</sub> | 13 | 13 | +1 |
| **Motixafortide** | Fba-RR(Nal)CY(Cit)K(dK)PYR(Cit)CR-NH<sub>2</sub> | 14 | 10 | +6 |

\* For RaPID-identified peptides, cyclization occurs between N-terminal ClAc-(dF) and the underlined Cys residue. For motixafortide, cyclization occurs Cys4 and Cys13. ClAc-(dF), chloroacetic-D-phenylalanine; Fba, 4-fluorobenzoyl; Nal, L-3-naphthylalanine; Cit, L-citrulline; dK, D-Lysine.

**Table S2: Potency of each identified peptide in signaling assays.**

| Receptor | CXCR4 | CXCR4 | CXCR4 | ACKR3 |
| --- | --- | --- | --- | --- |
| Ligand | Mini-Gα<sub>s/i</sub><br>(pIC<sub>50</sub>) | Gα<sub>i</sub> Activation<br>(pIC<sub>50</sub>) | β-arrestin2<br>(pIC<sub>50</sub>) | β-arrestin2<br>(pEC<sub>50</sub>) |
| **A** | < 5.00 | < 5.00 | < 5.00 | < 5.00 |
| **B** | < 5.00 | < 5.00 | < 5.00 | 7.62 |
| **C** | 6.47 | < 5.00 | < 5.00 | 6.08 |
| **D** | 6.25 | < 5.00 | < 5.00 | 5.71 |
| **E** | < 5.00 | < 5.00 | < 5.00 | < 5.00 |
| **F** | 6.22 | 6.18 | < 5.00 | 6.25 |
| **G** | 7.30 | < 5.00 | 6.38 | 7.00 |
| **H** | 6.83 | 5.58 | 6.66 | 7.64 |
| **I** | 7.64 | 6.69 | 6.97 | 3.08 |
| **K** | 7.54 | 5.92 | 7.19 | < 5.00 |
| **IT1t** | 7.35 | 6.29 | 8.10 | – |
| **VUF11207** | – | – | – | 8.10 |

**Table S3.** List of control chemokines in the chemokine receptor selectivity screening.

| *Chemokine*<br>*Receptor* | *Control*<br>*Chemokine* |
| --- | --- |
| CCR1 | CCL3 |
| CCR2B | CCL2 |
| CCR3 | CCL13 |
| CCR4 | CCL22 |
| CCR5 | CCL5 |
| CCR6 | CCL20 |
| CCR7 | CCL19 |
| CCR8 | CCL1 |
| CCR9 | CCL25 |
| CCR10 | CCL27 |
| CXCR1 | CXCL8 |
| CXCR2 | CXCL8 |
| CXCR3A | CXCL11 |
| CXCR3B | CXCL11 |
| CXCR4 | CXCL12 |
| CXCR5 | CXCL13 |
| CXCR6 | CXCL16 |
| XCR1 | XCL2 |
| CX3CR1 | CX3CL1 |
| ACKR2 | CCL5 |
| ACKR3 | CXCL12 |
| ACKR4 | CCL19 |

**Table S4:** <sup>1</sup>H assignment table of peptide I in water.

|  |  | <sup>1</sup>H δ (ppm) |  |  |  |  |  |  |  |  |  |  |  |  |
| --- | --- | --- | --- | --- | --- | --- | --- | --- | --- | --- | --- | --- | --- | --- |
| Res. # | Residue | NH | Ha | Hβ |  | Hγ | Hδ |  | Hε |  | Hζ |  | Hη | Other |
| 1 | Phe | 8.13 | 4.46 | 2.59 | 2.76 | – | 6.96 |  | 7.15 |  | 7.09 | – | 2.97* |  |
| 2 | Tyr | 8.32 | 4.25 | 2.43 | 2.71 | – | 6.68 | 6.55 |  |  | – |  | – |  |
| 3 | Ala | 8.05 | 4.10 | 1.11 |  | – | – |  | – |  | – | – | – |  |
| 4 | His | 8.18 | 4.26 | 3.00 | 3.06 | – | 8.39 | 7.10 | 7.08 | 8.37 |  | – | – |  |
| 5 | Arg | 8.18 | 4.02 | 1.60 | 1.65 | 1.38 | 2.97 |  |  |  | – | 7.01 | – |  |
| 6 | Leu | 7.70 | 4.14 | 1.39 | 1.46 | 1.39 | 0.69 | 0.72 | – |  |  | – | – |  |
| 7 | Arg | 7.86 | 3.85 | 1.55 |  | 1.38 | 2.97 |  |  |  | – | 6.99 | – |  |
| 8 | Arg | 7.69 | 4.08 | 1.46 | 1.51 | 1.27 | 2.92 |  |  |  | – | 6.93 | – |  |
| 9 | Trp | 7.95 | 4.49 | 3.09 | 3.17 | – | 7.03 | 10.03 | 7.05 | 6.99 | 7.25 | 7.45 | – |  |
| 10 | Leu | 7.85 | 4.27 | 1.35 | 1.42 | 1.24 | 0.65 |  |  | – | – | – | – |  |
| 11 | Ser | 8.07 | 4.22 | 3.70 |  | – | – |  | – |  | – | – | – |  |
| 12 | Cys | 8.08 | 4.29 | 2.47 | 2.57 | – | – |  | – |  | – | – | – |  |

\* CH<sub>2</sub> in thioether bond.
